# Two dynamic, N-terminal regions are required for function in Ribosomal RNA Adenine Dimethylase family members

**DOI:** 10.1101/2024.04.19.590347

**Authors:** Danielle A. McGaha, Alexandrea Collins, Luqman O. Ajisafe, Calvin C. Perdigao, Jordan L. Bondrowski, Karen Fetsch, Jack A. Dunkle

## Abstract

The Ribosomal RNA Adenine Dimethylase (RRAD) family of enzymes facilitate ribosome maturation in all organisms by dimethylating two nucleotides of small subunit rRNA. Prominent members of this family are the human DIMT1 and bacterial KsgA enzymes. A sub-group of RRAD enzymes, named erythromycin resistance methyltransferases (Erm) dimethylate a specific nucleotide in large subunit rRNA to confer antibiotic resistance. How these enzymes regulate methylation so that it only occurs on the specific substrate is not fully understood. While performing random mutagenesis on the catalytic domain of ErmE, we discovered that mutants in an N-terminal region of the protein that is disordered in the ErmE crystal structure are associated with a loss of antibiotic resistance. By subjecting site-directed mutants of ErmE and KsgA to phenotypic and in vitro assays we found that the N-terminal region is critical for activity in RRAD enzymes: the N-terminal basic region promotes rRNA binding and the conserved motif likely assists in juxtaposing the adenosine substrate and the SAM cofactor. Our results and emerging structural data suggest this dynamic, N-terminal region of RRAD enzymes becomes ordered upon rRNA binding forming a cap on the active site required for methylation.

## Introduction

In all organisms, RNA is post-transcriptionally modified in diverse ways, including methylation, thiolation, the conversion of uridines to pseudouridines and many other chemical marks (Boccaletto et al. 2018). Methylation is among the most frequent modifications of RNA occurring on tRNA and rRNA in all organisms and on mRNA in eukaryotes (Grosjean 2009; Machnicka et al. 2014; Roundtree et al. 2017). Remarkably, a limited number of protein topologies are required to carry-out methylation of diverse RNA substrates. The two most prominent protein topologies are the SPOUT (SpoU-TrmD) fold and the Rossmann-like fold (Czerwoniec et al. 2009). The SPOUT fold possesses five parallel beta strands with the topology ↑2-↑1-↑4-↑3-↑5 and the distinctive feature of a C-terminal α-helix which forms a trefoil knot by threading between a loop linking β-strand 3 and β-strand 4 (Strassler et al. 2022). Methyltransferases with the Rossmann-like fold are also known as Class I methyltransferases and possess seven β-strands with the topology ↑3-↑2-↑1-↑4-↑5-↓7-↑6 (Schubert et al. 2003). β-strands 1-3 of Class I RNA methyltransferases serve as a platform for structural elements that bind the S-adenosylmethionine cofactor while β-strands 4-7 play the same role for RNA binding (Fig. 1A.)

**Figure 1.**
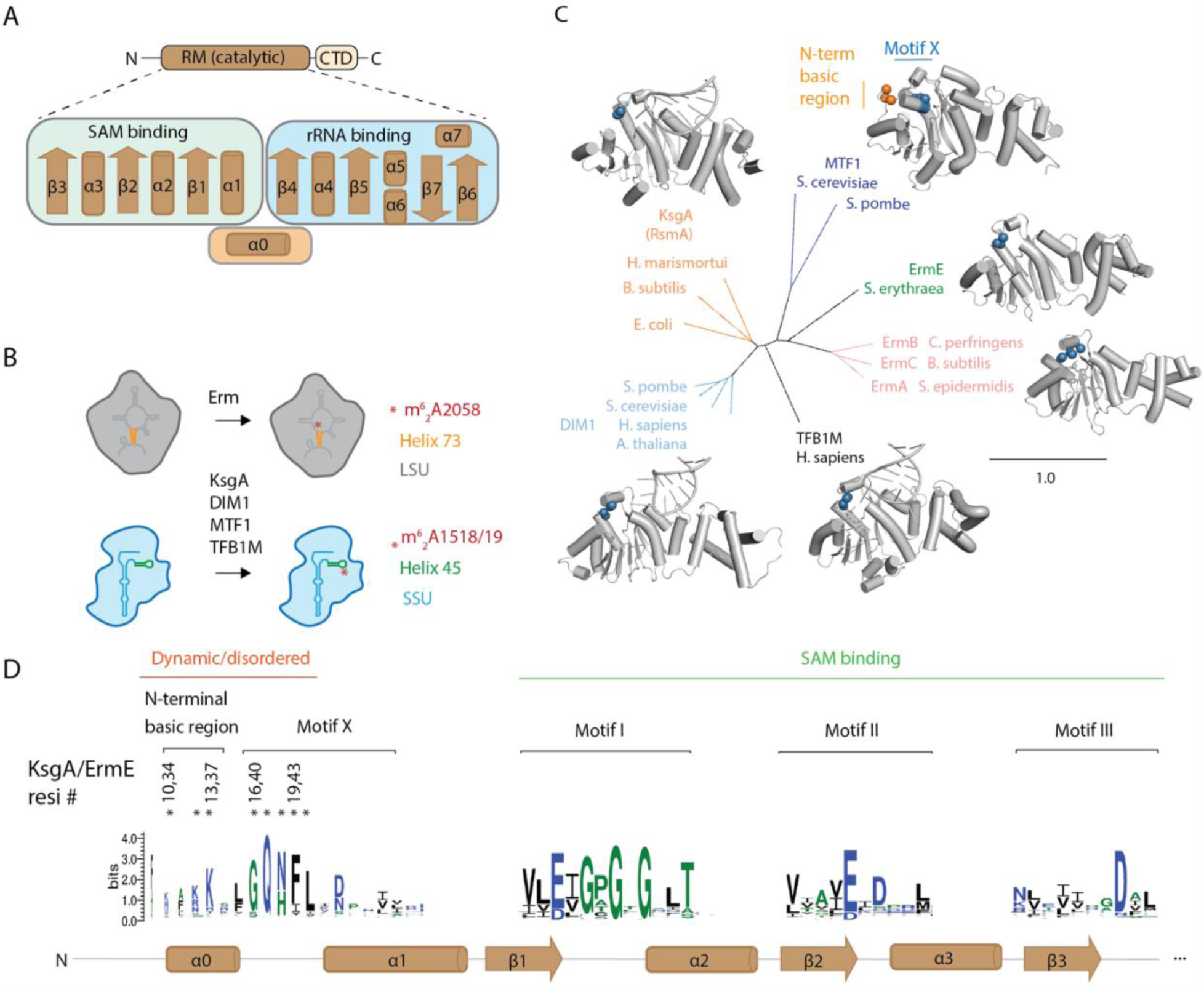
The structure and activity of Ribosomal RNA Adenine Dimethylase (RRAD) family members. (A) RRAD enzymes consist of a Rossmann-like methyltransferase (RM) catalytic domain and a C-terminal domain. The secondary structure of the RM domain is shown indicating α1-β3 are predominantly involved in *S*-adenosylmethionine (SAM) while β4-β7 are involved in rRNA binding. (B) RRAD family members dimethylate rRNA of either the large ribosomal subunit (LSU) or small ribosomal subunit (SSU). (C) Experimentally determined structures are available for each RRAD family member in some cases bound to rRNA (MTF1, PDB code 1i4w; ErmE, 6nvm; ErmC, 1qao; TFB1M, 6aax; DIM1, 7wtm; KsgA, 7o5h). An N-terminal basic region (orange) is disordered in these structures (except for MTF1) while the adjacent Motif X (blue) is partially ordered. (D) The conservation of RRAD family members at key motifs throughout the α0-β3 region of the protein family is shown as a sequence logo. Residues in the N-terminal basic region and Motif X investigated by site-directed mutagenesis in this study are marked with *.

Class I methyltransferases participate in critical functions in both eukaryotes and prokaryotes. For example, METTL1 in humans carries out m^7^G modification of tRNA (Li et al. 2023) (Ruiz-Arroyo et al. 2023). In metazoans METTL3, along with its METTL14 partner, deposits m^6^A in mRNAs to regulate their stability and the dynamic status of m^6^A marks in mRNA has been implicated in many disease states (Dominissini et al. 2012; Meyer et al. 2012; Murakami and Jaffrey 2022). In metazoans, METTL16 has been found to introduce m^6^A into the U6 snRNA and an intronic region of the MAT2A transcript regulating its stability and therefore the levels of its protein product, a SAM synthetase (Pendleton et al. 2017). In prokaryotes, deposition of m^7^G at position 1405 or m^1^A at position 1408 of 16S rRNA provides bacteria resistance to aminoglycoside antibiotics (Jeremia et al. 2023). The dimethylation, m^6^_2_A2058, on 23S rRNA provides multi-drug antibiotic resistance to macrolide, lincosamide and streptogramin B antibiotics (MLS_B_ phenotype) (Fyfe et al. 2016). Interestingly, while the overall architecture of these enzymes is diverse and the type of methylation varies, the Class I methyltransferase catalytic domain is shared by all (Dunkle et al. 2014; Sledz and Jinek 2016; Wang et al. 2016; Doxtader et al. 2018; Srinivas et al. 2023).

The Ribosomal RNA Adenine Dimethylase (RRAD) family comprise enzymes with sequence and structural homology that utilize a Class I catalytic domain to dimethylate adenosine residues on small ribosomal subunit RNA or large ribosomal subunit RNA (Mistry et al. 2021) (Figs. 1B, 1C). KsgA deposits m^6^ A at positions 1518 and 1519 (*E. coli* numbering) of 16 S rRNA in bacteria during ribosome biogenesis (Fig. 1B). The current model for KsgA function is that it binds a conformation of the 30S ribosome that is off-pathway for maturation and this binding event promotes remodeling of the 30S to an on-pathway conformation (Connolly et al. 2008; Sun et al. 2023). Since ribosomes are able to assemble in Δ*ksgA* cells, methylation is not required for ribosome biogenesis but rather promotes KsgA dissociation from the ribosome once the remodeling event has occurred (Connolly et al. 2008). DIMT1 (DIM1) performs the same methylations on the cytoplasmic ribosomes of eukaryotes (Lafontaine et al. 1994; Zorbas et al. 2015; Shen et al. 2020), whereas TFB1M in humans methylates the 12S rRNA of mitochondrial ribosomes (Metodiev et al. 2009) (Liu et al. 2019). Erythromycin resistance methyltransferases, such as ErmE or ErmC, dimethylate a single position, A2058, in 23S rRNA of bacteria during large ribosomal subunit biogenesis to promote antibiotic resistance (Fyfe et al. 2016) (Fig. 1B). ErmE is found in *S. erythraea*, a soil bacterium that biosynthesizes the protein synthesis inhibitor erythromycin and it protects the bacterium from this toxic molecule (Skinner et al. 1983). ErmC is found on mobile genetic elements that circulate widely among Gram-positive bacteria such as *B. subtilis* and *S. aureus*. ErmC is an significant contributor to multi-drug antibiotic resistance in *S. aureus*.

In all cases RRAD proteins are thought to bind *S*-adenosylmethionine (SAM) and the nucleotide targeted for methylation in a similar manner using conserved motifs (Fig. 1D). RRAD proteins contain conserved motifs observed in other 6-methyladenosine methyltransferases (Schubert et al. 2003). These include Motif I, a Gly-rich region that sits adjacent to the kink between the ribose and methionyl moieties of SAM and has shape complementarity to it. Motif II possesses an acidic residue that hydrogen bonds to the ribose moiety of SAM. Motif IV possesses an aromatic residue that forms π-stacking interactions to the target Ade residue. The importance of these interactions has been validated by numerous functional studies (Farrow et al. 2002; Maravic et al. 2003; O’Farrell et al. 2012; Rowe et al. 2020; Shen et al. 2020; Goh et al. 2022; Sharkey et al. 2022).

We asked whether there may be unappreciated elements of the catalytic domain of RRAD proteins required for function. To answer this question, we performed random mutagenesis on the catalytic domain of ErmE and screened the resulting cells for an erythromycin sensitive phenotype. Surprisingly, the screen indicated residues in an unstructured N-terminal region were associated with an erythromycin sensitive phenotype. Site-directed mutants of ErmE were generated and characterized phenotypically and by in vitro biochemistry. These data indicated the existence of two unappreciated regions in ErmE that are critical for function: an N-terminal basic region and Motif X (Fig. 1D). We next investigated whether these regions were functionally important in other Erm proteins using the ErmC model system and in RRAD proteins, broadly, using the small subunit rRNA methyltransferase KsgA as a model. We produced evidence that the N-terminal basic region and Motif X are also required for function in these methyltransferases indicating they are a characteristic of RRAD family members generally. A synthesis of our biochemical data and available structural data suggests the N-terminal region undergoes a disordered to ordered transition when basic residues recognize the enzyme’s specific rRNA substrate and the ordered conformation contributes key interactions to the active site.

## Results

We previously identified multiple residues within the catalytic domain of ErmE that are required for function (Rowe et al. 2020; Sharkey et al. 2022). Residues were target for site-directed mutagenesis based on sequence alignment to ErmC or because a Rosetta docking of ErmE to its rRNA substrate indicated they were likely to contact rRNA (Fig. 2A). We wondered whether there may be residues in ErmE critical for function that had not yet been identified by a model driven approach. Therefore, we performed random mutagenesis of the catalytic domain of ErmE by error-prone PCR (Figs. 2A, S1). The plasmid containing the resulting library was used to transform *E. coli* cells. Cells containing the ErmE plasmid were selected by their growth on an ampicillin plate, then were transferred to an erythromycin plate with the expectation that loss of catalytic activity in ErmE would result in an erythromycin sensitive phenotype. From a pool of 182 transformants we identified 146 colonies that were erythromycin sensitive. We sequenced these transformants, and all contained multiple mutations, as expected for our implementation of error-prone PCR. We plotted the frequency that each amino acid was altered in an erythromycin sensitive colony (Fig. 2B). The mutations at S88 and A96 were frequently observed which could be rationalized based on their proximity to the *S*-adenosylmethionine (SAM) binding site. Random mutagenesis of ErmB has previously been reported and also observed mutations in the SAM pocket associated with an erythromycin sensitive phenotype (Farrow et al. 2002). Additionally, R163 is a position that closely approaches rRNA in our computational model and whose site-directed mutants are erythromycin sensitive (Sharkey et al. 2022). Unexpectedly, we also saw P20 and R47 mutations often occurred among our erythromycin sensitive colonies which are both located in the N-terminus which is not ordered in the crystal structure of ErmE (Stsiapanava and Selmer 2019).

**Figure 2.**
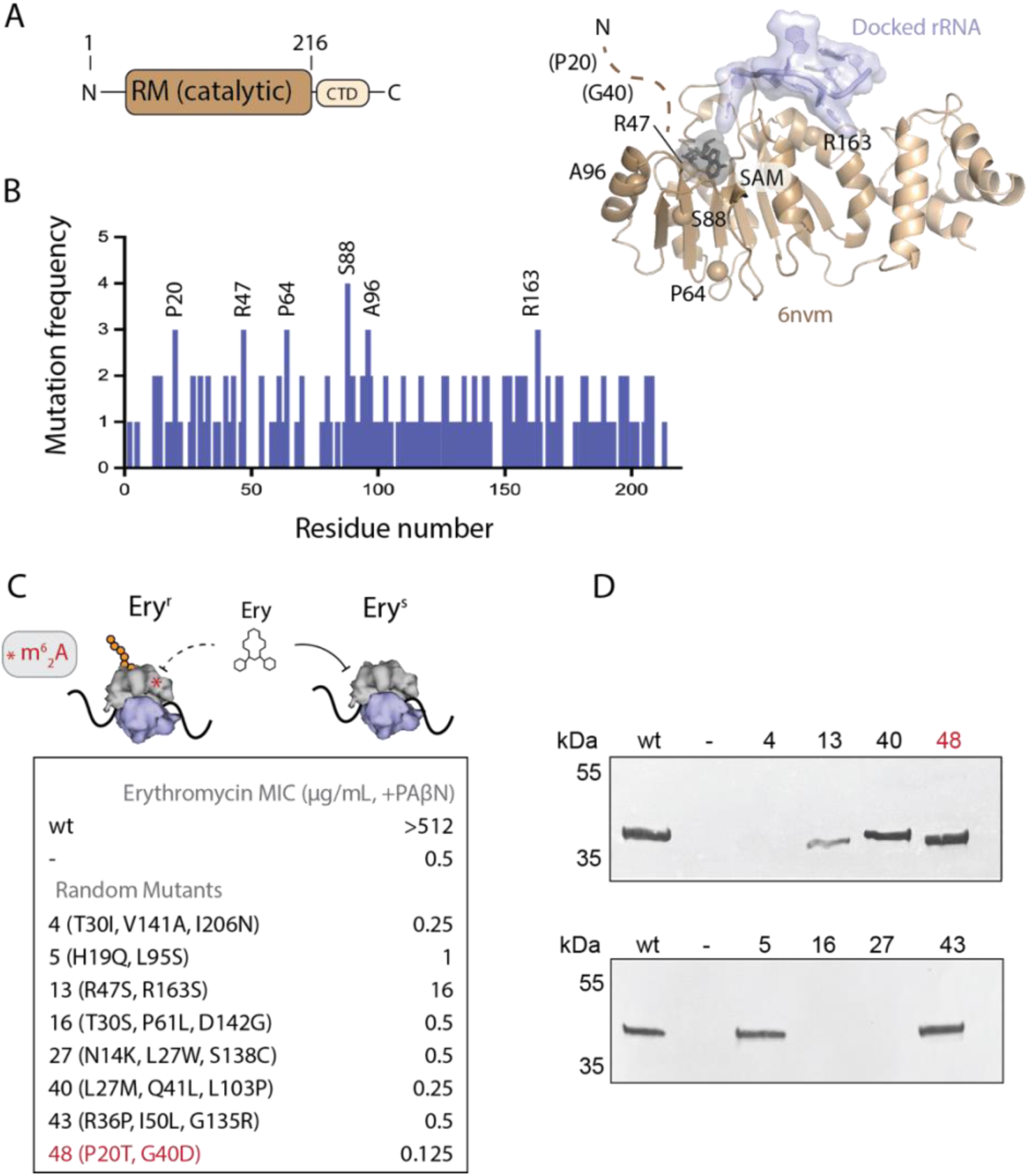
Random mutagenesis reveals mutations in the dynamic, N-terminal region of ErmE are associated with an erythromycin sensitive phenotype. (A) The crystal structure of the RRAD family member ErmE lacks the dynamic, N-terminal region. Docking of SAM onto the crystal structure and rRNA via Rosetta modeling gives the approximate positions of the substrates. The residues most frequently associated with an erythromycin sensitive phenotype in random mutagenesis screens are shown as spheres. Residues P20 and G40 were not present in the coordinate model. (*B*) Random mutagenesis of the catalytic, Rossmann-like methyltransferase (RM) domain of ermE was performed. A histogram shows how often specific ermE residues were mutated among clones that were erythromycin sensitive. (*C*) Minimal Inhibitory Concentrations (MIC) for erythromycin were measured in *E. coli* cells expressing *ermE* mutants in the presence of the antibiotic adjuvant phenyl-arginyl-beta-napthylamide (PAβΝ). Random mutagenesis produced variants with 2-4 mutations per clone. (C) Western blotting of cell lysates was used to detect whether *ermE* mutants produced a soluble and stable protein.

### Mutants of the basic region and Motif X are associated with an erythromycin sensitive phenotype

To scrutinize the observation that mutations in the dynamic N-terminal region may contribute to an erythromycin sensitive phenotype, we performed a microdilution assay on a selection of these mutants. The assay confirmed that the selected colonies possessed very low minimal inhibitory concentrations (MIC) for erythromycin (Fig. 2C). We performed western blotting of these mutants to determine which random mutants may owe their phenotype to a destabilization of the ErmE structure and which may be explained by a loss of ErmE catalytic activity. Four out the eight mutants assayed produced a signal comparable to wildtype in a blot of the supernatant of lysed *E. coli* cells suggesting the mutants did not destabilize ErmE (Fig. 2D). Interestingly, one mutant, the P20T G40D double mutant, was solely composed of mutations to the dynamic, N-terminal region of ErmE suggesting it may play a previously unappreciated role in ErmE function.

Since the N-terminal region of ErmE is not ordered in the crystal structure we inspected an AlphaFold2 model of ErmE to hypothesize how this region may contribute to function (Jumper et al. 2021). AlphaFold2 predicts the presence of an alpha helix enriched in basic residues followed by a loop region with sequence conservation among many DNA and RNA methyltransferases that has been denoted as Motif X (Malone et al. 1995)(Figs. 1D and 3A). The N-terminal basic region and Motif X are adjacent to the ErmE active site. We hypothesized that the basic region, while disordered in the apo structure of ErmE, may become ordered upon RNA binding. It may contribute to the binding affinity of ErmE for RNA or may influence the conformation of Motif X in a manner that promotes SAM binding or positioning of the target A2058 nucleotide in the active site. To assess the importance of the basic region and Motif X for ErmE function, we generated site-directed mutants of the two regions and scored the resulting phenotype of cells bearing these mutants in the presence of erythromycin (Fig. 3B). We altered basic Arg to the acidic Glu and polar or hydrophobic residues to Ala. Three site-directed mutants (R36P, R47S and G40D) were constructed due to observations from random mutagenesis. Most, but not all, mutants displayed a dramatic reduction in MIC value. To dissect whether the erythromycin sensitive phenotypes were derived from a loss of ErmE catalytic activity or destabilization of the protein we again performed western blots. Within the basic region, site-directed mutants of R34, R36 and R37 were associated with erythromycin sensitivity yet produced levels of protein similar to wild type suggesting these mutants are defective in catalysis (Figs. 3B and 3C). Random mutagenesis had indicated R47 may play an important functional role in ErmE, but the R47S mutant was not erythromycin sensitive (Fig. 3B). Mutants G40D, N42A and F43A within Motif X were associated with erythromycin sensitivity but had levels comparable to wild type by western blotting suggesting a catalytic defect (Figs. 3B and 3C). Taken together, our phenotypic experiments identified multiple residues within both the basic region and Motif X that appear to be crucial for catalysis.

**Figure 3.**
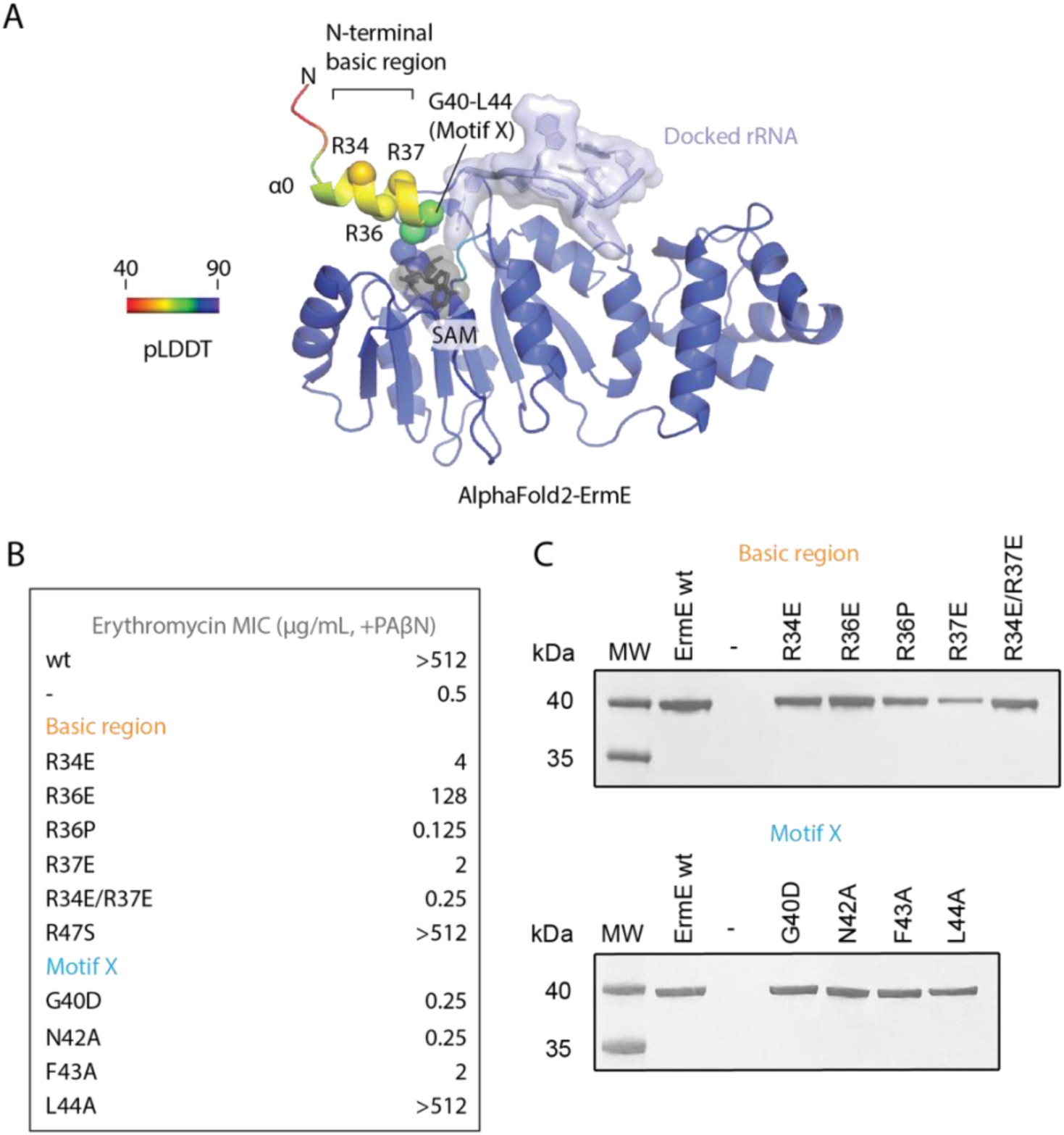
Site-directed mutants in the N-terminal basic region and Motif X are associated with altered phenotypes in ErmE. (*A*) An AlphaFold2 model of ErmE suggests some of the dynamic N-terminal residues form α-helix 0. Superposition of the AlphaFold2 ErmE model onto the model given in Figure 2A indicates α0 forms one wall of the active site. (*B*) MIC values for erythromycin were measured for *E. coli* cells expressing *ermE* site-directed mutants in the basic region and Motif X. (*C*) Western blotting of cell lysates was used to detect whether *ermE* site-directed mutants produced a soluble and stable protein.

To further test the hypothesis that the basic region and Motif X contribute to ErmE function we performed in vitro methylation reactions. The reactions were carried out with wild type and site-directed mutants of ErmE, a synthetic RNA that forms a hairpin mimicking helix 73 of 23S rRNA and ^3^H-SAM as methyl donor (Fig. 4A, Fig. S2). The experiments were performed under single turnover conditions, that is with excess enzyme, and with SAM limiting. Mutants reporting on the contribution of the basic region, R34E/R37E and R36P, showed very low levels of methylation compared to wild type (Fig. 4A). However, R36E possessed an intermediate defect. Mutants reporting on Motif X, G40D and N42A, had low levels of methylation while the F43A mutant had an intermediate level of methylation (Fig. 4A). These results strongly suggest both the basic region and Motif X are critical for ErmE function and are consistent with the results from the phenotypic assays. The two site-directed mutants with intermediate levels of methylation, R36E and F43A, also had intermediate MIC values of 128 μg/mL and 2 μg/mL erythromycin, respectively in the phenotypic assays, demonstrating a qualitative correlation between methyltransferase activity and erythromycin resistance.

**Figure 4.**
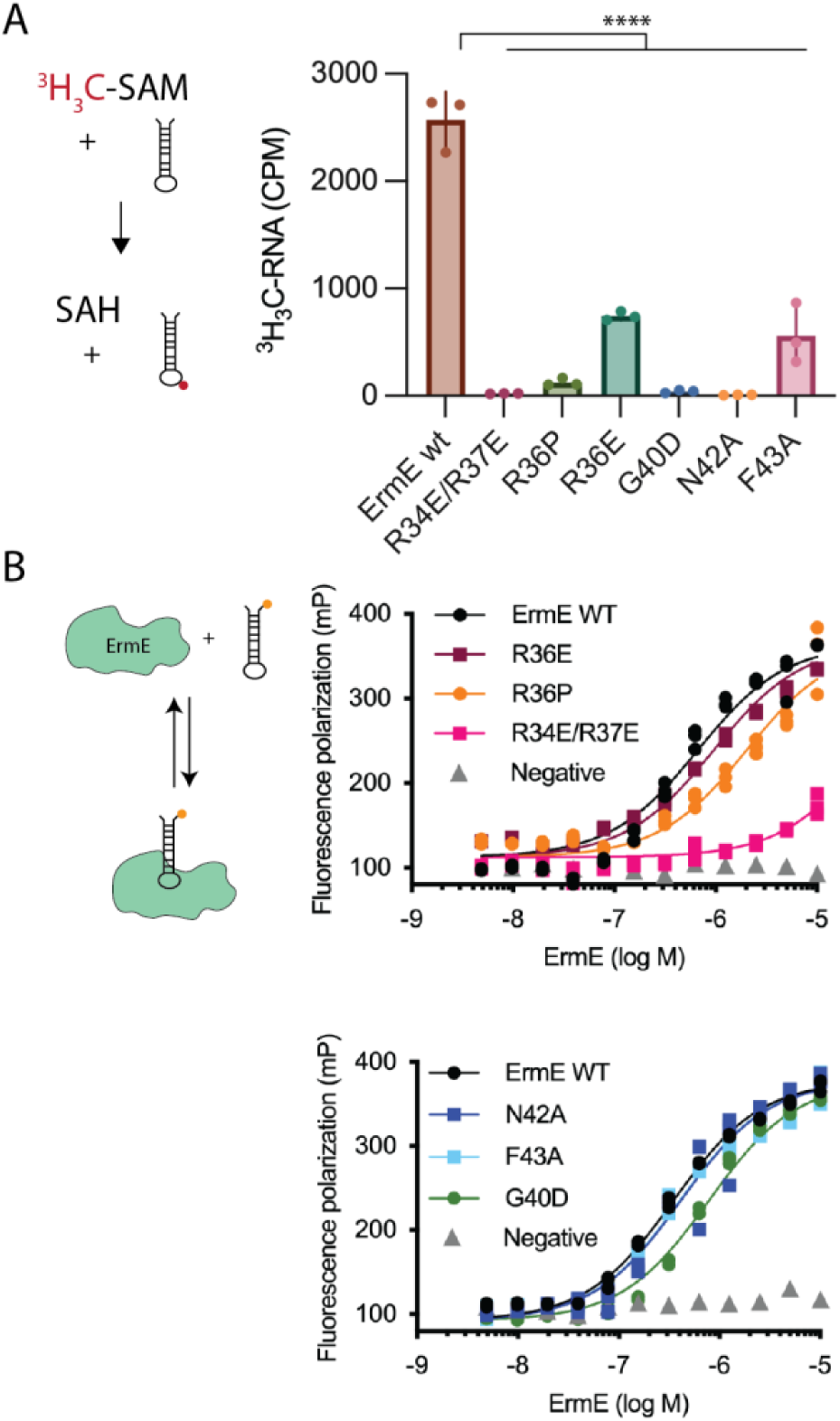
The N-terminal basic patch and Motif X contribute to rRNA methylation and the basic patch contributes to rRNA binding. (A) Methylation reactions under single-turnover conditions were conducted using site-directed mutants in the N-terminal basic region or Motif X of ErmE. (B) Affinity binding of ErmE site-directed mutants to an analog of helix 73 of 23S rRNA was measured by fluorescence polarization. A cartoon depicting the assays is given. The fluorescein label on the 48-nt rRNA analog is indicated by an orange circle. ****, p < 0.001.

### Identifying the mechanistic contributions of the basic region and Motif X to catalysis

To begin to dissect how the basic region and Motif X contribute to rRNA methylation by ErmE, we performed a saturation binding assay to test how mutants affected affinity for rRNA. Purified site-directed mutants of ErmE were titrated against a helix-73 rRNA analog possessing a fluorescent label and binding was read by fluorescence polarization (Fig. 4B). ErmE wild type was used as a positive control for binding and pepsin, a protein not known to interact with RNA, was used as a negative control. The basic region mutant R36E had a binding affinity similar to wild type. R36P possessed a modest defect in RNA binding while an R34E/R37E double mutant displayed a major defect. The Motif X mutants N42A and F43A possessed binding curves similar to wild type, but G40D had a modest defect in binding. Taken together these data indicate that Motif X does not make a major contribution to RNA binding, but the basic region does. Alteration of a single residue in the basic region does not lead to a major defect in RNA binding but alteration of multiple residues, as in R34E/R37E does.

With, the RNA affinity binding data in hand, we reasoned we could combine these data with structural information and a kinetic analysis of ErmE and its mutants to determine the most likely contribution of the basic region and Motif X to ErmE catalysis. We sought to distinguish between contributions to rRNA binding, SAM binding or chemistry, such as positioning of the nucleophilic N6 of adenosine adjacent to the sulfonium of SAM. The structure of ErmC bound to SAM, the apo structure of ErmE, our Rosetta model of ErmE bound to a fragment of rRNA and the crystal structure of RRAD family member TFB1M bound to RNA informed our analysis (Schluckebier et al. 1999; Liu et al. 2019; Stsiapanava and Selmer 2019; Sharkey et al. 2022). While Motif X is not fully ordered in these structures, the portion that is ordered, for example N36, F37 and L38 in TFB1M, suggest it could contribute to SAM binding affinity or positioning of the target Ade or SAM for chemistry. The RNA binding affinity reported above shows it does not make a major contribution to rRNA binding. The possibilities for the basic region are more complex. While our data show that it makes a contribution to RNA binding, that interaction with RNA could influence the adjacent Motif X residues to affect SAM binding or chemistry or both.

To distinguish between the possible ways the basic region and Motif X could contribute to catalysis, we followed methylation over time, under conditions of limiting SAM and as a function of ErmE concentration (Fig. 5A). We extracted observed rate constants (k_obs_ min^-1^) for these reactions and plotted them versus ErmE concentration fitting the data to a hyperbola to produce K_1/2,SAM_ and k_chem_, which reflects the rate limiting step prior to product formation. The K_1/2,SAM_ parameter should approximate K_d,SAM_ since our reaction is under rapid equilibrium conditions (Hou and Masuda 2015) (Fig. 5B, Table 1). We chose the mutants R36E and F43A for this analysis because they retained enough methylation activity to produce a signal that could be fit to our kinetic model.

**Figure 5.**
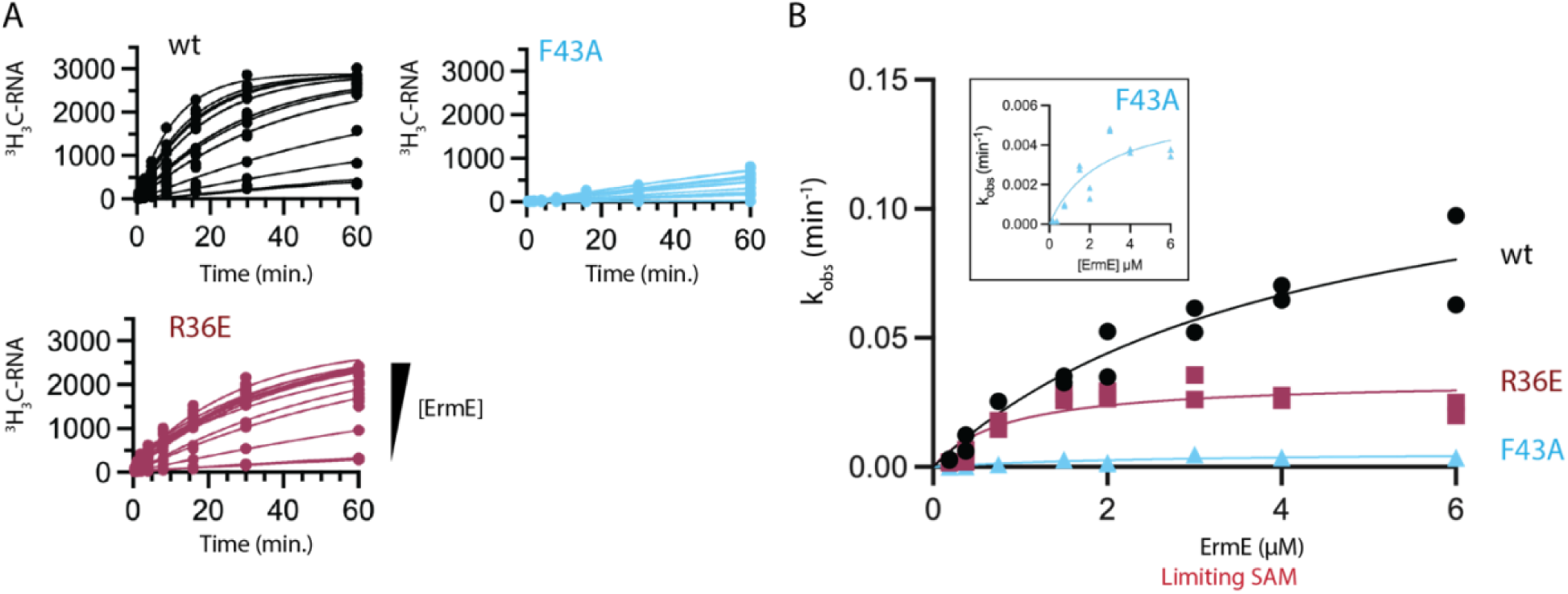
Single turnover kinetics. (A) Methylated RNA versus time plots for increasing concentration of wild type (wt) ErmE and site-directed mutants. (B) Methylation rates (k_obs_) are plotted versus ErmE concentration under limiting SAM and excess RNA conditions and fit to a model for single-turnover kinetics to extract k_chem_ and K_1/2,SAM_ values. An inset is shown of the F43 data with a y-axis range of 0 to 0.006 min^-1^.

**Table 1.** Single turnover kinetics and RNA affinity binding parameters.

| | $k_{\text{chem}}$ ($\text{min}^{-1}$) | $K_{1/2, \text{SAM}}$ ($\mu\text{M}$) | $K_d, \text{RNA}$ ($\mu\text{M}$) |
| --- | --- | --- | --- |
| wt | 0.14 (0.092-0.23) | 4.2 (2.1-10) | 0.66 (0.55-0.79) |
| R36E | 0.033 (0.027-0.044) | 0.83 (0.38-1.8) | 2.0 (1.6-2.4) |
| F43A | 0.0061 (0.0040-0.013) | 2.6 (0.94-9.6) | 0.40 (0.35-0.47) |

Parentheses indicate 95% confidence interval.

R36E possesses a four-fold decrease in k_chem_ compared to wild type (0.033 min^-1^ versus 0.14 min^-1^) but possesses a K_1/2,SAM_ = 0.83 μM which is slightly better than wild type and indicates there is no defect in SAM binding (Table 1). R36E does bind RNA with slightly lower affinity than wild type (2.0 μM versus 0.66 μM) but since the kinetics assay is performed with [R36E] in excess of K_d_ the binding defect does not explain the 4-fold lower k_chem_. One possible explanation of the data is that the basic region binding to rRNA affects the structure in a way that enhances catalysis. F43A possesses a 23-fold decrease in k_chem_ compared to wild type (0.0061 min^-1^ versus 0.14 min^-1^) indicating a major defect in catalysis, yet possesses a K_1/2_,_SAM_ = 2.6 μM similar to wild type. Since we have shown above that Motif X mutants do not possess a major binding defect for RNA, the synthesis of the available data suggests how the basic region and Motif X affect the mechanism of ErmE. Binding of rRNA by the basic region enhances the affinity of ErmE for rRNA and subtly affects the structure of the neighboring Motif X in a manner that promotes catalysis. Since the residues comprising Motif X are not suspected of acid-base chemistry, it likely promotes catalysis by optimally positioning the N6 of the target Ade adjacent to the labile methyl group or SAM.

### The basic region and Motif X contribute to rRNA methylation by RRAD family members

Sequence alignments of ErmE with ErmC, other erythromycin resistance methyltransferases and other RRAD family members, suggests the presence of an N-terminal basic region and Motif X is a general feature of these proteins (Figs. 1D, S3). Additionally, comparison of the AlphaFold2 structures of ErmE and ErmC reinforces this (Figs. 3A, 6A). We generated site-directed mutants of ErmC within the basic region and Motif X, both single and double mutants, and assayed cells containing them for erythromycin sensitivity. As a positive control we assayed cells containing wild type ErmC and as a negative control, cells containing an empty vector. The K4E single mutant possessed a reduced MIC value indicating erythromycin sensitivity and the K4E/K7E double mutant had a further reduced MIC value consistent with the basic region in ErmC playing an important functional role (Fig. 6B). Single mutants within Motif X did not possess erythromycin sensitivity but double mutants did (Fig. 6B). While the erythromycin sensitivity phenotype associated with these ErmC mutants was not as pronounced as in ErmE, the data suggest that both the basic region and Motif X also play an important role in ErmC function.

**Figure 6.**
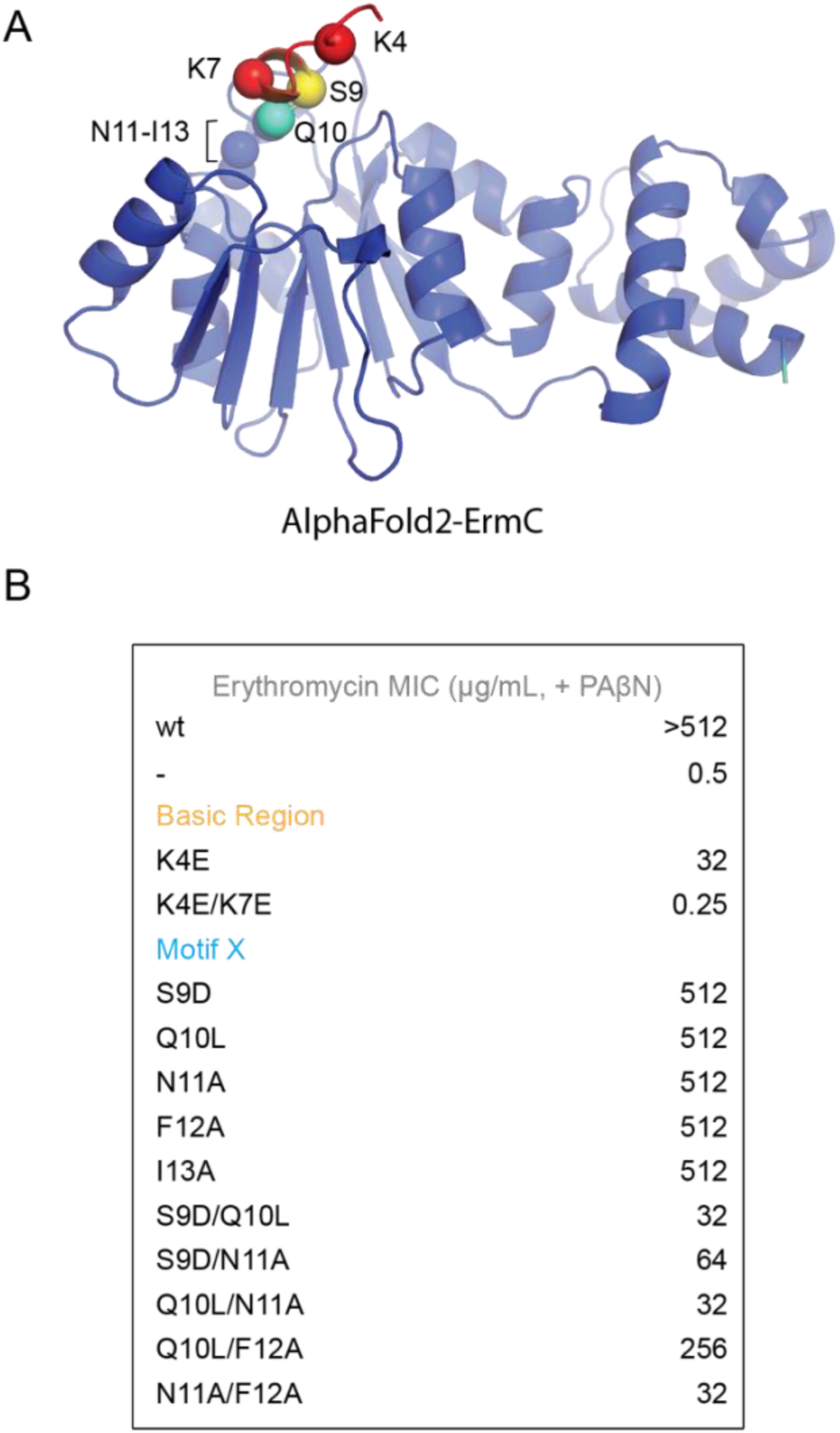
Site-directed mutants in the N-terminal basic region and Motif X are associated with altered phenotypes in ErmC. (A) An AlphaFold2 model of ErmC demonstrating the predicted location of basic region residues K4 and K7 and motif X residues S9-I13. (B) MIC values for erythromycin were measured for *E. coli* cells expressing *ermC* site-directed mutants in the basic region and motif X

The Erm proteins are the only members of the RRAD family that methylate large ribosomal subunit rRNA; all other family members methylate small ribosomal subunit rRNA (Fig. 1B). KsgA (RsmA) in bacteria dimethylates two adenosine residues in helix 45 of 16S rRNA as part of its role in promoting 30S ribosome maturation. To determine whether the N-terminal basic region and Motif X play a role in the function of the small ribosomal subunit rRNA methyltransferases we used the bacterial KsgA protein as a model system.

We developed a phenotypic assay that reports on KsgA methylation of 16S rRNA by utilizing the aminoglycoside antibiotic kasugamycin. This molecule inhibits initiation of bacterial protein synthesis by binding to a pocket on the ribosome that overlaps with the path of mRNA between the E and P-sites of the 30S subunit (Poldermans et al. 1979; Schuwirth et al. 2006) (Fig. 7A). This pocket is adjacent to the sites methylated by KsgA, m^6^ A1518 and m^6^ A1519. Loss of methylation on A1519, for example in Δ*ksgA* cells, leads to partial resistance to kasugamycin (Helser et al. 1972). It appears the loss of methylation affects the conformation of residues adjacent to A1519 that directly interact with the antibiotic (Fig. 7A). Notably, this phenomenon wherein methylation causes antibiotic sensitivity, works in the opposite manner of ErmE or ErmC where methylation causes antibiotic resistance. Site-directed mutants of KsgA in the basic region and Motif X were constructed and the ability of cells bearing these mutants to grow in the presence of kasugamycin was assayed (Figs. 7B, 7C). Three controls were employed to validate the assay worked in the intended manner, that is, growth of cells which efficiently form m^6^ A1519 on 16S rRNA is inhibited by kasugamycin, but cells which do not efficiently methylate this residue are kasugamycin resistant. First, *E. coli* K12 BW25113 Δ*ksgA* from the Keio collection, labeled as “Vector” in Figure 7C, were transformed with empty vector (Baba et al. 2006). As expected, these cells grew in the presence of kasugamycin, although growth was slower than in the no kasugamycin scenario confirming that loss of m^6^ A1519 leads to partial resistance. Second, we hypothesized that the Y116A mutant of KsgA would be deficient in methylation and found this was the case (Figs. 8A, 8B). Many Class I RNA methyltransferases possess an aromatic residue in Motif IV that π-stacks with the substrate nucleobase (Dunkle et al. 2014; Liu et al. 2019). This Motif IV residue has been shown to be strictly required for rRNA methylation by Erm enzymes and a multiple sequence alignment shows that Y116A is the Motif IV aromatic residue in KsgA (Maravic et al. 2003) (Rowe et al. 2020) (Fig. S3). Cells bearing Y116A possessed a low level of inhibition by kasugamycin confirming that the growth defect in the presence of kasugamycin is substantially dependent on rRNA methylation (Fig. 7C). Our final control was to measure the growth of the ΔksgA cells complemented with wild type ksgA from a plasmid. These cells were strongly inhibited by kasugamycin, as expected (Fig. 7C).

**Figure 7.**
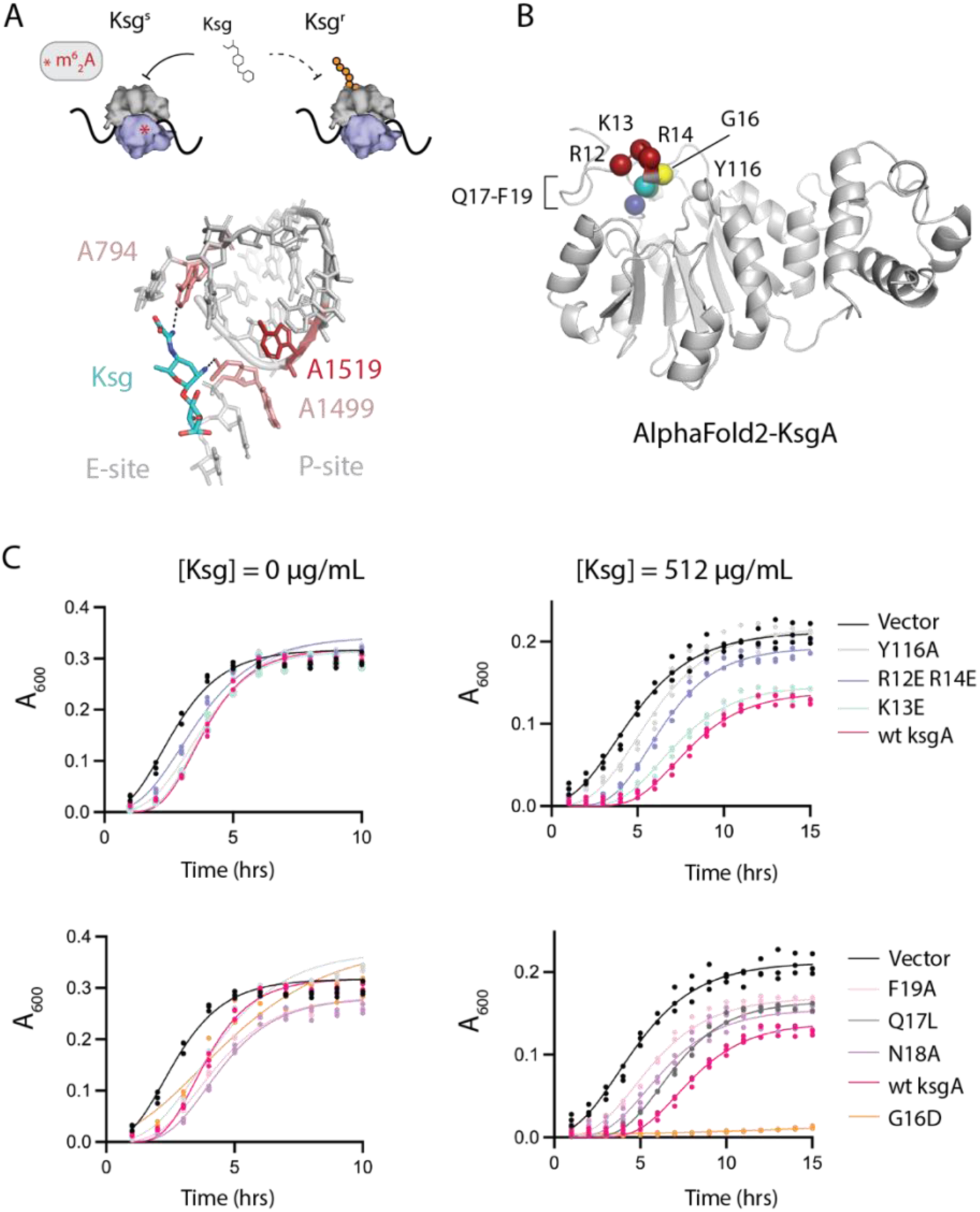
The N-terminal basic patch and Motif X of RRAD family member, KsgA, contribute to function in vivo. (A) Kasugamycin (Ksg) inhibits protein synthesis by binding to ribosomes that possess the m^6^ A1519 modification on 16S rRNA. Methylation is indicated by the red asterisk. The structure of Ksg bound to the *E. coli* ribosome is shown (PDB code 1vs5). A loss of methylation on 16S rRNA residue A1519 affects the local conformation leading to partial Ksg resistance. (B) An AlphaFold2 model of *E. coli* KsgA shows the predicted location of the basic region R12-R14 and Motif X residues G16-F19. (C) Growth curves of *E. coli* Δ*ksgA* in the absence or presence of kasugamycin. The cells are transformed with wild type wt *ksgA*, a site-directed mutant of *ksgA* or empty vector (Vector). Y116A is included as a control because it is defective in methylation.

**Figure 8.**
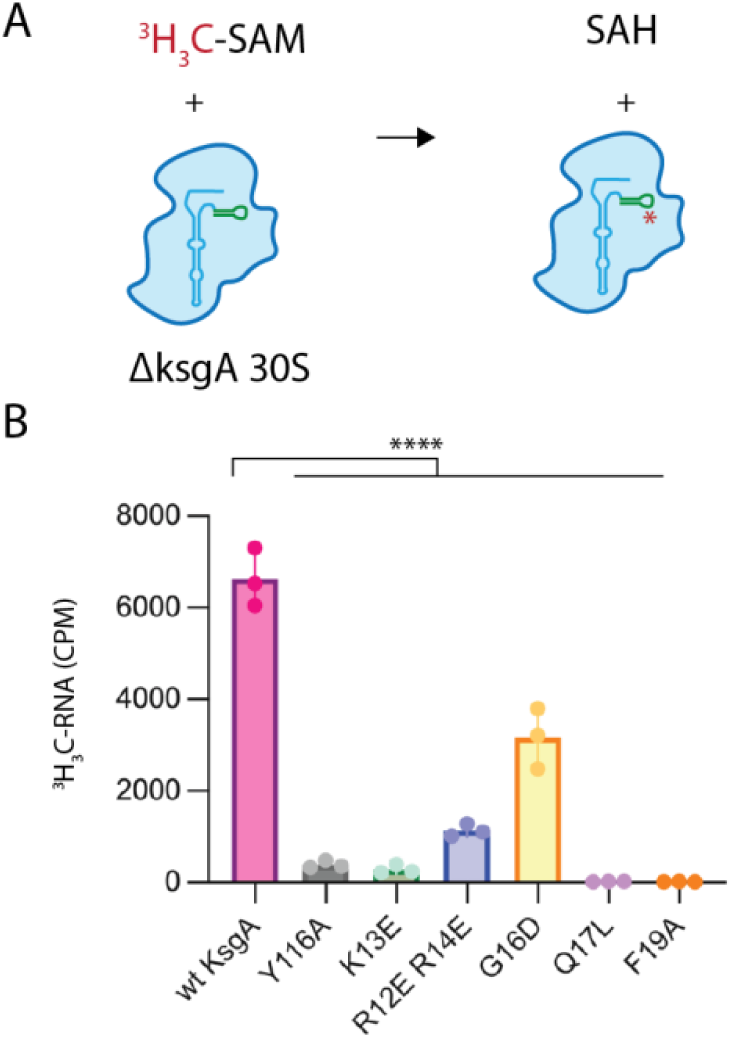
The N-terminal basic region and Motif X of KsgA contribute to 16S rRNA methylation in vitro. (A) KsgA catalyzed methylation of 16S rRNA in vitro was measured using 30S ribosomes devoid of the m^6^ A1518 and m^6^ A1519 (Δ*ksgA* strain) and radiolabeled SAM cofactor. (B) Wt KsgA robustly methylates 30S ribosomes in vitro. The Y116A active site mutant is methylation deficient and serves as a negative control. ****, p < 0.001.

Having validated that our phenotypic assay produces results that correlate with the methylation activity of KsgA, we subjected site-directed mutants in the basic region and Motif X to the assay. While basic region mutant K13E in the presence of kasugamycin grew at a rate similar to wild type, the R12E/R14E double mutant grew better than wild type consistent with decreased methylation activity. Motif X mutants Q17L, N18A and F19A all grew better than wild type consistent with decreased methylation. Recombinant expression of G16D produced an unexpected result strongly inhibiting cell growth. This mutant likely interferes with ribosome biogenesis but an investigation of this is beyond the scope of our work. Circular dichroism data indicate this mutant may be partially unfolded (Fig. S4). Taken together, these results are consistent with the hypothesis that the basic region and Motif X contribute to methylation activity in all RRAD family members.

To further test the hypothesis that the basic region and Motif X are crucial for KsgA activity, we purified selected variants and performed in vitro methylation assays (Figs. 8A, 8B, S4). The assays employed 30S ribosomal subunits purified from Δ*ksgA* cells as substrate and ^3^H-SAM (Fig. S5). Methylation of 30S subunits by wild type KsgA was performed as a positive control and reactions containing the Y116A variant of KsgA were used as a negative control (Fig. 8B). Basic region mutants K13E and R12E/R14E had markedly decreased methylation activity (Fig. 8B). The fact that K13E has a phenotype similar to wild type KsgA yet performs in vitro methylation poorly was unexpected. However, this may be explained by the fact that in vivo methylation occurs on immature 30S subunits as a step in ribosome maturation and our in vitro methylation assays use fully formed 30S subunits which are likely poorer substrates for KsgA. Motif X mutants Q17L and F19A both exhibit levels of methylation that are similar to our negative control, while G16D possessed about one-third as much methylation activity as wt. In sum, the in vitro methylation results are consistent with what we observed in the phenotypic assays and demonstrate the basic region and Motif X contribution to methylation by KsgA.

## Discussion

We have provided functional evidence that two, unappreciated regions of RRAD family enzymes are critical for their function. We demonstrated the importance of the N-terminal basic region and Motif X using in vivo assays of site-directed mutants of ErmE, ErmC and KsgA. We performed methylation assays with purified site-directed mutants of ErmE and KsgA to verify that mutants with a phenotype suggesting loss-of-function did in fact have perturbed methylation. Using ErmE as a model system we performed RNA affinity binding assays and kinetics assays to determine the mechanistic contribution of these regions to rRNA methylation. We found that the N-terminal basic region contributes to rRNA binding. Motif X mutants bind rRNA normally and our kinetics investigation of the Motif X mutant, F43A, indicated it has a K_1/2, SAM_ similar to wild type indicating it binds SAM normally. This leads to the question: why are Motif X mutants defective in methylation? Given its structural location, adjacent to the sulfonium of SAM and the adenosine ring of the target nucleotide, and the fact that it doesn’t possess ionizable residues typically associated with chemical steps in catalysis, Motif X most likely is involved in optimally positioning the substrate to react with the sulfonium of SAM.

Considering our biochemical data alongside existing structural data it seems likely that the N-terminal basic region and Motif X are typically dynamic in the absence of rRNA, but binding of the basic-region to its specific rRNA substrate, orders it and the neighboring Motif X to promote catalysis (Fig. 9A). In support of this idea, most crystal structures of RRAD family enzymes in the absence of RNA have a disordered N-terminal basic region and a partially ordered Motif X. Three examples of this are the apo structures of ErmC, ErmE and *E. coli* KsgA (PDB codes 1qao, 6nvm and 1qyr) (Schluckebier et al. 1999) (O’Farrell et al. 2004; Stsiapanava and Selmer 2019). Interesting exceptions to this are the *B. subtilis* KsgA structures reported in PDB code 6ifs (Bhujbalrao and Anand 2019). Two copies of KsgA are present in the asymmetric unit with chain B possessing a partially ordered basic region and a gap of ten disordered residues, containing part of Motif X, before the chain becomes ordered again. Chain A in this structure has both regions fully ordered. Cryo-EM structures of *B. subtilis* KsgA bound to the 30S ribosome were recently solved capturing five distinct KsgA bound states (Singh et al. 2022). The state referred to as K5 by the authors has a disordered N-terminal basic region and Motif X, as well as disordered helix 44 of 16S rRNA, but in the K1-k4 state both KsgA regions are ordered as is helix 44 (Figs. 9B, 9C). In recent structures of the human RRAD enzyme TFB1M bound to an oligo mimicking rRNA, the N-terminal basic region and part of Motif X are disordered but in cryo-EM structures of TFB1M bound to the small subunit of the mitochondrial ribosome these regions are ordered (PDB codes 6aax and 8csp) (Liu et al. 2019; Harper et al. 2023). However, in recent cryo-EM structures of *E. coli* KsgA bound to immature 30S ribosomes from the Ortega and Davis groups, a different mechanism from that in Figure 9A was proposed. It was argued that the N-terminal region of KsgA interacts with helix 24 (h24) of 16S rRNA and that this interaction keeps h24 from occluding the KsgA active site (Sun et al. 2023).

**Figure 9.**
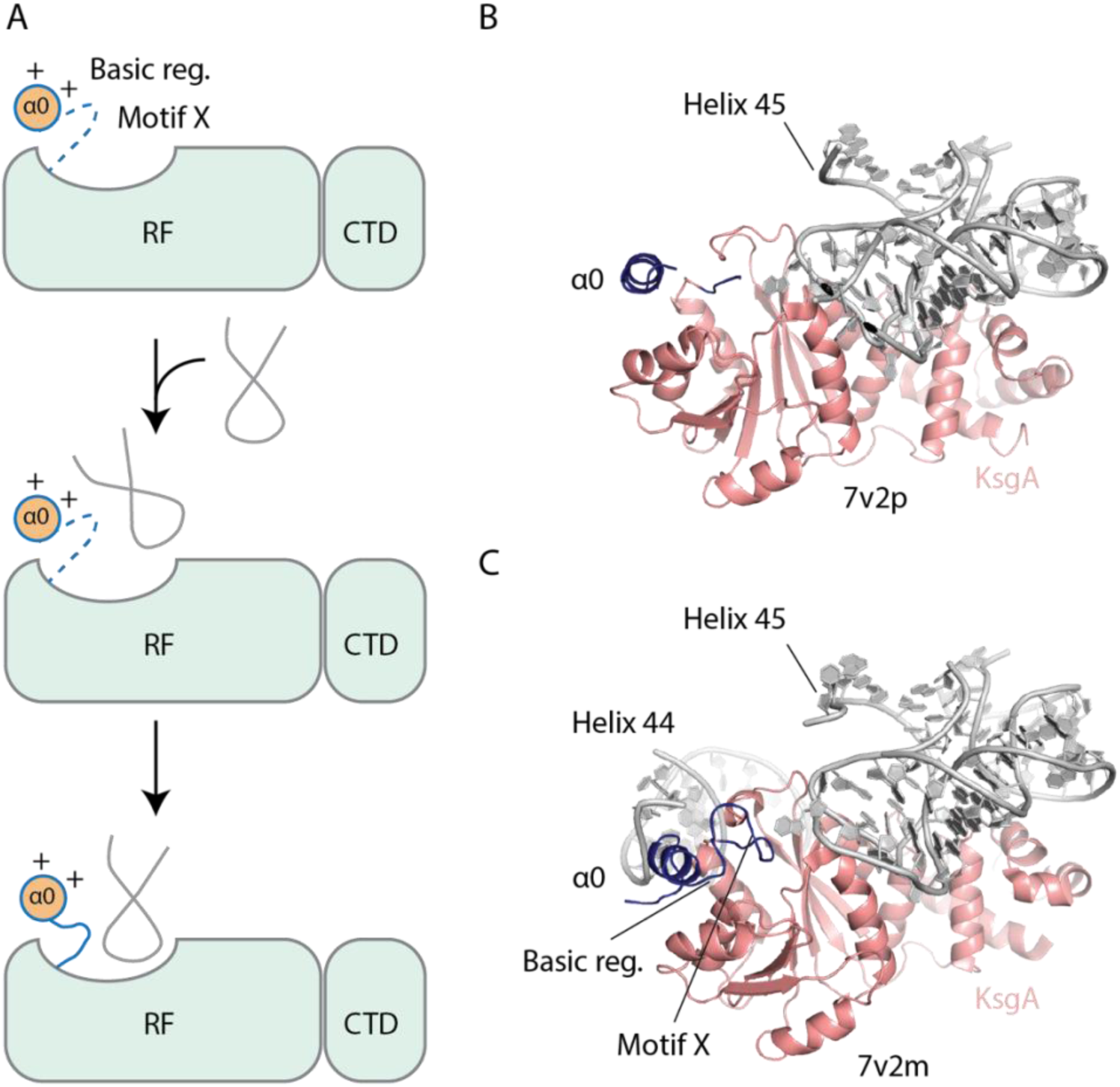
A model for the contribution of the basic region and Motif X to rRNA methylation. (A) In the absence of rRNA, Motif X is disordered. The electrostatic interaction of the basic region with rRNA promotes an initial encounter complex that then matures into a structure with an ordered Motif X contributing to the active site of the RRAD methyltransferase. (B) In the KsgA-30S ribosome structure reported in PDB code 7v2p, helix 44 of 16S rRNA is disordered as is the basic region and Motif X of KsgA, but in (C) the structure reported in 7v2m helix 44 is ordered along with the basic region and Motif X. The basic region is positioned to make electrostatic contacts to helix 44 of 16S rRNA.

The phenomenon of a dynamic N-terminal region that is critical for function has been reported recently for two other Rossmann-like, RNA methyltransferases indicating this may be a common mechanism among these enzymes. Human METTL1, with the assistance of partner protein WDR4, installs m^7^G46 on multiple tRNAs, a modification that regulates tRNA stability and is associated with disease states (Shaheen et al. 2015; Dai et al. 2021; Orellana et al. 2021). The N-terminal region possesses sequence conservation and is the location of a phosphorylation site, at residue S27, that regulates its activity, yet in structures of METTL1-WDR4 in the absence of its tRNA substrate this region is disordered (Li et al. 2023; Ruiz-Arroyo et al. 2023). In cryo-EM structures of METTL1-WDR4 bound to tRNA, the N-terminal region is ordered and forms interactions adjacent to the G46 target nucleotide thought to be important for positioning it for catalysis (Li et al. 2023; Ruiz-Arroyo et al. 2023). Functional assays by the same authors validate the importance of the N-terminal region: the site-directed mutant R24A and phosphomimetic mutants of S27 have dramatically reduced methylation. Rossmann-like methyltransferase TlyA methylates positions in both 16S and 23S rRNA and the presence of these methylations facilitates binding of the anti-tuberculosis antibiotics, capreomycin and viomycin, to the ribosome similar to the situation with KsgA and kasugamycin (Johansen et al. 2006). Full-length TlyA is recalcitrant to crystallization, but TlyA-CTD readily crystallizes (Witek et al. 2017). Two crystal forms of truncated TlyA have different positions of the peptide that links the NTD and CTD which extrapolate to different spatial arrangements of the full-length protein. The cryo-EM structure of TlyA bound to the 50S ribosome confirms a critical role for the NTD in rRNA recognition (Laughlin et al. 2022). Collectively, the available structural and functional data suggest that recognition of rRNA by the NTD of TlyA affects the RAWV linker peptide whose conformation affects SAM binding and methylation.

Two interesting future directions for understanding how the N-terminal region of RRAD proteins contributes to catalysis are: what are the specific interactions between Motif X and the substrates that are driving catalysis and is the N-terminal region a modular appendage that controls RNA substrate specificity? The former question arises because the KsgA and TFB1M ribosome-bound structures are determined by cryo-EM at moderate resolution and therefore don’t produce certainty regarding side-chain positions. Additionally, the side chain-substrate interactions are likely dynamic across the reaction coordinate and therefore functional studies and structures of multiple reaction intermediates will be necessary to truly describe them. The latter question pre-supposes that since the fold of Class I RNA methyltransferases is conserved, the variable appendages must be providing the RNA substrate specificity. Evidence supporting this model was recently reported by Bhujbalrao and colleagues who showed that KsgA/ErmC chimeras that included the N-terminus could alter substrate specificity (Bhujbalrao and Anand 2019). The many roles RNA methyltransferases play in health and disease warrants a thorough understanding of their structures and mechanisms.

## Materials and Methods

### Random mutagenesis of ermE

Random mutants were created using pBAD/Myc-His-ermE as template DNA, a construct previously reported (Rowe et al. 2020). Briefly, *ermE* codon-optimized for E. coli was inserted into pBAD/Myc His A (Invitrogen) resulting in an open reading frame encoding the sequence given by Uniprot IDP07287 except that the 82 carboxy-terminal residues, which encode a low complexity Gly-rich segment, are removed and the MYC and hexa-histidine tags are added to the carboxy terminus. GeneMorph II Random Mutagenesis Kit (Agilent Technologies) was used according to manufacturer instructions except that to achieve a low mutation frequency, 100 ng of target DNA was used as well as 20 PCR cycles for Mutant Megaprimer synthesis. Transformants were plated on LB-ampicillin (100 μg/mL) plates that were coated with 250 μL of 0.2% w/v L-arabinose. To determine the phenotype of each random mutant an agar-based selection assay was performed. Transformants were transferred to a gridded 0.2% w/v L-arabinose coated LB-ampicillin (100 μg/mL) plate as well as a gridded 0.2% w/v L-arabinose coated LB plate containing 512 μg/mL of erythromycin. The plates were incubated at 37°C overnight. Colonies that did not grow on the LB-erythromycin plate were then collected from the LB-ampicillin plate for plasmid purification and sequenced via Sanger sequencing (Eurofins).

### Erythromycin MIC assay

Microdilution MIC assays were performed similarly to a previous protocol with minor modifications (Sharkey et al. 2022). Colonies were resuspended in 300 mM NaCl and diluted to an A_600_ of 0.1. A 2-fold serial dilution was performed in a sterile, 96-well assay block of Mueller-Hinton broth containing: erythromycin ranging from 0 μg/mL to 512 μg/mL, 0.02% w/v L-arabinose, and 2.5 mg/mL PAβN in a 200 μL final volume. A volume of 2 μL of the cell suspension was transferred to the assay block at each of the concentrations. The assay block was incubated at 37°C for 16 h. A plate reader was used to assess growth, with an A_600_ value of 0.01 greater than the background signal indicating growth. The lowest concentration of erythromycin able to prevent cell growth was recorded.

### Westen blotting

Western blotting was performed similarly to a previous protocol with minor modifications (Rowe et al. 2020). Cells were harvested from the microdilution assay block from wells containing 0 μg/mL erythromycin media. The cells were harvested by centrifugation, resuspended in Buffer E (50 mM Tris HCl pH 7.5, 250 mM NaCl, 1 mM DTT, 2% v/v glycerol) with lysozyme (0.1 mg/mL), and incubated on ice for 30 min. Cells were then lysed with two freeze/thaw cycles in liquid nitrogen followed by centrifugation to pellet cellular debris. The supernatant was subjected to SDS-PAGE on a 4–20% gel, with wild type and pBAD (empty vector) as controls, then transferred to a nitrocellulose membrane. The membrane was blocked for 1 h in 3% w/v BSA in TBST, washed with TBST three times, and incubated for 1 h in 1:1250 anti-myc antibody at room temperature. The membrane was washed again with TBST and incubated in 1:2500 anti-mouse HRP conjugated antibody for 1 h. The membrane blots were developed with 1-Solution TMB and imaged.

### ErmE Site-directed mutagenesis

Site-directed mutagenesis of pBAD/Myc-His-ermE was previously described (Rowe et al. 2020). Site-directed mutants were generated in one of two ways as indicated in Supplemental Table S1 along with the oligonucleotide sequences used.

### ErmE purification

To purify the ErmE variants for in vitro assays, *E. coli* TOP10 cells containing either pBAD-*ermE* or site-directed mutants were cultured in LB media at 37°C until reaching an optical density of ∼0.6 at A_600_. Recombinant expression was induced by adding 0.02% w/v L-arabinose, and cell growth was contined at 37°C for 4 hours. Cells were harvested via centrifugation, and the pellet was resuspended in Buffer E, along with 15 mM imidazole and 0.1% v/v Triton X-100. Sonication was employed to disrupt the cells, and after centrifugation at 4800g for 30 minutes, the clarified lysate was applied to HisPur immobilized Ni^2+^ resin. The resin was washed with 15 column volumes of Buffer E supplemented with 15 mM imidazole. Protein elution was accomplished through a stepwise gradient of increasing imidazole concentrations (125 mM, 250 mM, and 500 mM) over 15 column volumes. Fractions were subjected to SDS-PAGE analysis to evaluate purity. Further purification of ErmE variants was conducted via size-exclusion chromatography using an S75 (Sephadex) column. Fractions containing monomeric ErmE were pooled and concentrated using a Pall centrifugal device with a 10,000 kDa MWCO membrane. Glycerol was then added to a final concentration of 10% v/v, and aliquots were flash-frozen for storage until required.

### RNA affinity binding

RNA binding affinity was assessed using fluorescence polarization as previously reported (Sharkey et al. 2022). Briefly a synthetic 48-nucleotide RNA mimicking helix 73 of 23S rRNA (V48), purified via PAGE and labeled with fluorescein at the 5′ end (obtained from Horizon Discovery), served as the substrate (Vester et al. 1998). Protein serial dilutions, ranging from 20 μM to 10 nM were combined with fluorescein labeled RNA and incubated for 2 hours at room temperature prior to fluorescence detection on a Bio-Tek Synergy H1 plate reader. The assay was performed in 0.5x Buffer E. A titration of the protein pepsin served as a negative control. Data from three replicates were fitted to the equation: y = B_max_∗x/(K_d_ + x) + y_0_ in GraphPad Prism.

### ErmE endpoint methylation assay

Methylation assays were conducted under single-turnover conditions following previously described methods with slight modifications (Rowe et al. 2020). All reactions were conducted in Buffer E with ErmE at 10 μM, RNA at 10 μM and ^3^H-SAM was present at 0.05 μM (55-85 Ci/mmol). The synthetic oligonucleotide V48 served as the RNA substrate (Vester et al. 1998; Rowe et al. 2020). After 60 min., 2.5 μL of the reaction mixture were withdrawn and quenched by diluting into 47.5 μL of 0.1 mg/mL salmon sperm DNA, followed by addition of 200 μL of 10% TCA. The precipitated RNA was collected via vacuum filtration using a Millipore Multiscreen GF 96-well plate, washed with 10% TCA and ethanol, and subsequently dried. Scintillation counting was performed on a MicroBeta2 instrument using PerkinElmer Betaplate scintillation fluid.

### ErmE single turnover kinetics

Single turnover kinetic assays were performed similarly to limiting SAM methylation assays described above except that ErmE or site-directed mutant was titrated from 6.0 to 0.2 μM and timepoints between 0 and 60 minutes were collected. Product versus time curves (Fig. 5A) were fitted to the expression y = (y_max_ - y_0_)*(1 - e^-kobs*x^) + y_0_ using GraphPad Prism. The k_obs_ for each reaction was plotted versus ErmE (μM). These curves were fit to the expression: k_obs_ = k_chem_* x/(K_1/2_ + x), where x =[ErmE], using GraphPad Prism.

### Construction of pBAD-ksgA and site-directed mutagenesis

Insertion of *E. coli ksgA* into pBAD/Myc-His was performed in the following manner. PCR was used to amplify the *ksgA* ORF from *E. coli* TOP10 genomic DNA. A second PCR reaction was used to add homology arms to pBAD/Myc-His. Finaly, an NEB HiFi assembly reaction was performed with the product of the second PCR reaction and linearized vector. The primers used in the assembly are given in Table S1.

### Growth kinetics in the presence of kasugamycin

Keio Collection *E. coli* K-12 BW25113 Δ*ksgA* cells cells were purchased from Horizon Discovery (Baba et al. 2006). *E. coli* K-12 BW25113 Δ*ksgA* harboring pBAD-ksgA WT or site-directed mutants were grown in LB media containing 0.02% w/v L-arabinose at 37°C for 16 h. Cells were then diluted to 0.2 A_600_. A serial dilution was performed in a sterile, 96-well assay block of Mueller-Hinton broth containing kasugamycin ranging from 0 mg/mL to 512 mg/mL, 2% w/v L-arabinose, and 20 mg/mL PAβN with 200 μL final volume. A volume of 10 μL of the cell suspension was transferred to the assay block at each of the concentrations. The assay block was incubated at 37°C for 16 h. A plate reader was used to assess growth every hour.

### 30S ribosomal subunit purification

*E. coli* K-12 BW25113 Δ*ksgA* was used to inoculate 10 mLs of LB which was grown overnight at 37°C. This culture was used to inoculate 1000 mL of LB which was grown at 37°C until it reached A_600_ ≃ 0.6. The culture was placed on ice for one hour and then the cells were collected by centrifugation. The cells were resuspended in Buffer A, 20 mM HEPES pH 7.5, 100 mM NH_4_Cl, 10 mM Mg(OAc)_2_ and 6 mM 2-mercaptoethanol. The cells were lysed by sonication and DNase I (RNase-free) was added. Centrifugation was used to remove cell debris and the supernatant was concentrated on a 100 kDa MWCO filtration device. The concentrated supernatant was layered onto a sucrose cushion made of Buffer A with 30% w/v sucrose. The ribosomes were pelleted by ultracentrifugation at 32k RPM for 22 hours. The pellet was dissolved in Buffer B, 20 mM HEPES pH 7.5, 50 mM NH_4_Cl, 1 mM Mg(OAc)_2_ and 6 mM 2-mercaptoethanol and layered onto a sucrose gradient composed of 10%-40% sucrose w/v in Buffer B. Ultracentrifugation of the gradient was performed in a SW32Ti rotor for 10 hr at 32k RPM. Fractions were eluted from the gradient using a syringe pump loaded with heavy sucrose and collected on a Biocomp fraction collection system with a Triax flow cell.

### KsgA methylation assay

Recombinant expression and purification of KsgA and its variants were performed in the same manner as the ErmE expression and purification described above. Methylation of 30S ribosomal subunits was performed in Buffer E (described above) diluted 1:1 with ultrapure water. 30S subunits at 10 μM and ^3^H-SAM at 0.1 μM final concentration were incubated with 10 μM KsgA at 37°C for 60 minutes. Reaction mixture was aspirated at various time intervals, quenched with 10% TCA and prepared for scintillation counting as described above for ErmE methylation assays.

## Acknowledgements

This work was supported by National Institute of General Medical Sciences of the National Institutes of Health under Award Number R35GM142966.

