## Supporting Information for "Two dynamic, N-terminal regions are required for function in Ribosomal RNA Adenine Dimethylase family members"

**Running title**: Two N-terminal regions required for RRAD function

**Keywords**: rRNA, methyltransferase, RNA modification, ribosome biogenesis, antibiotic resistance

**Supporting Information**

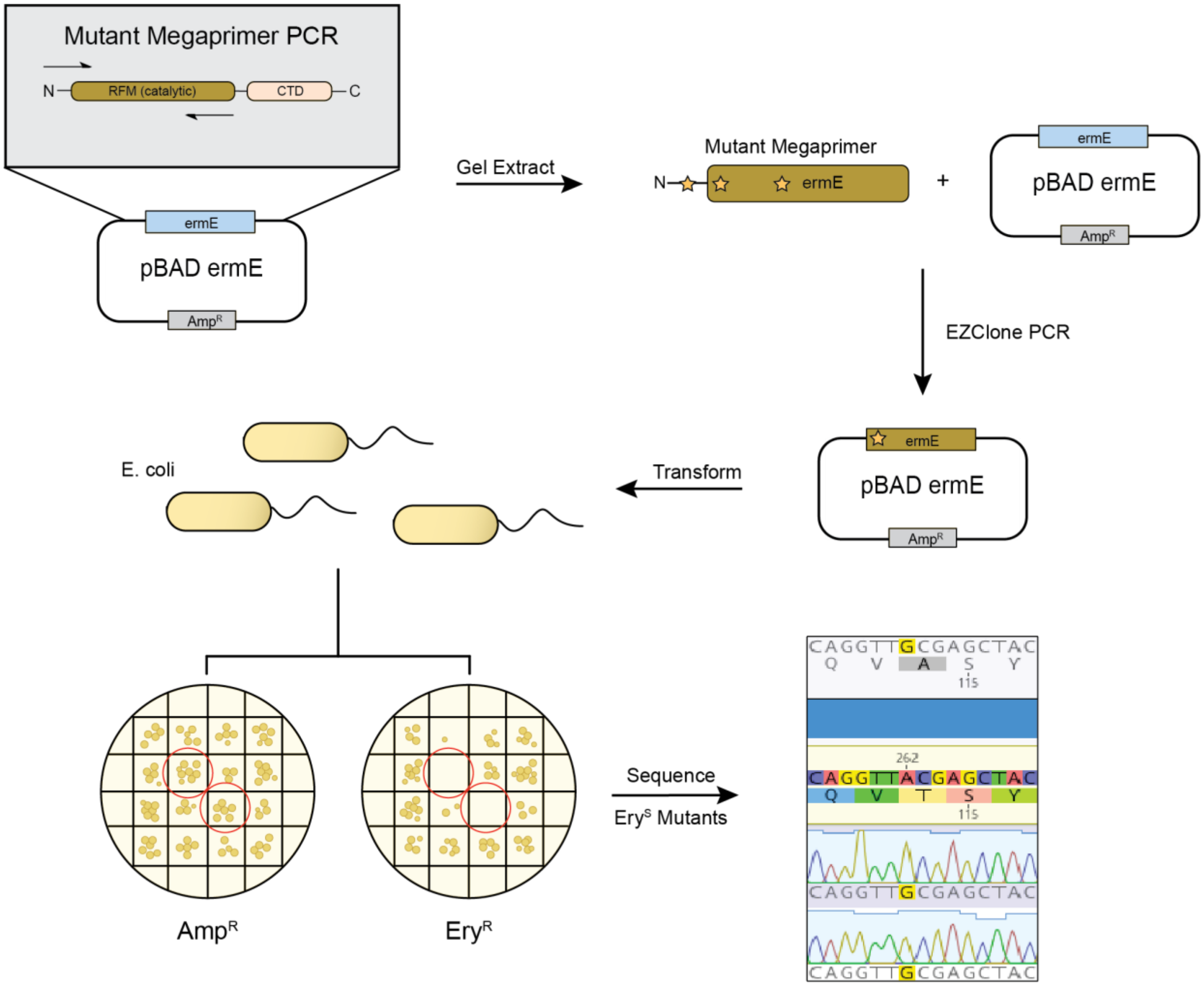


**Figure S1. A description of the random mutagenesis workflow**.

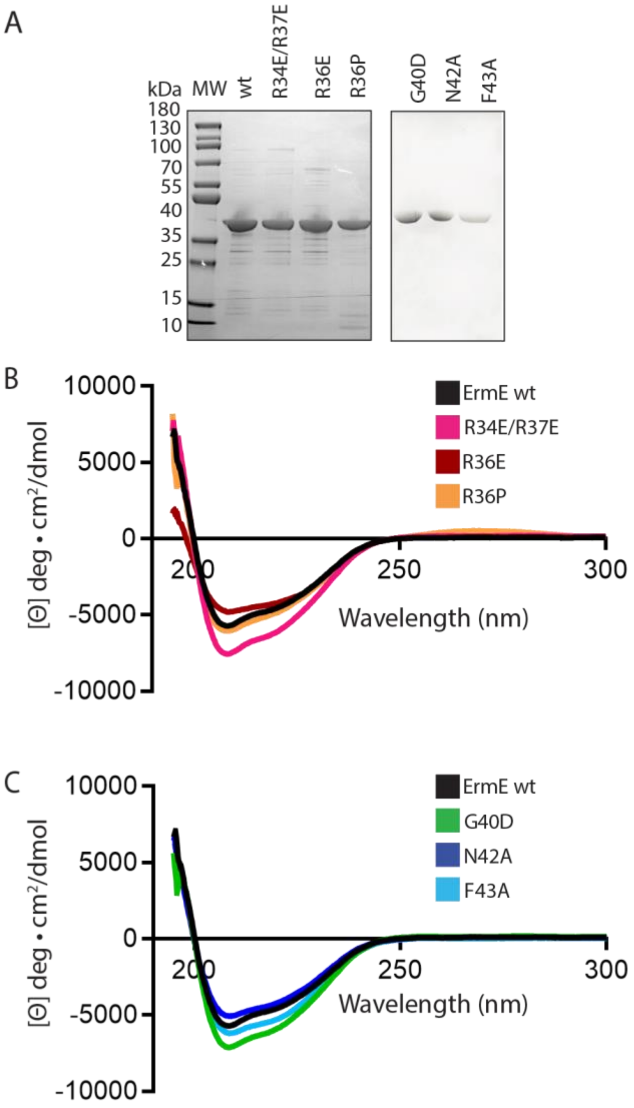


**Figure S2. Selected site-directed mutants of ErmE maintain native structure.** (A) SDS-PAGE analysis of wt ErmE and site-directed mutants is shown. (B) Circular dichroism spectroscopy of site-directed mutants within the N-terminal basic patch. (C) Circular dichroism spectroscopy of site-directed mutants within motif X.

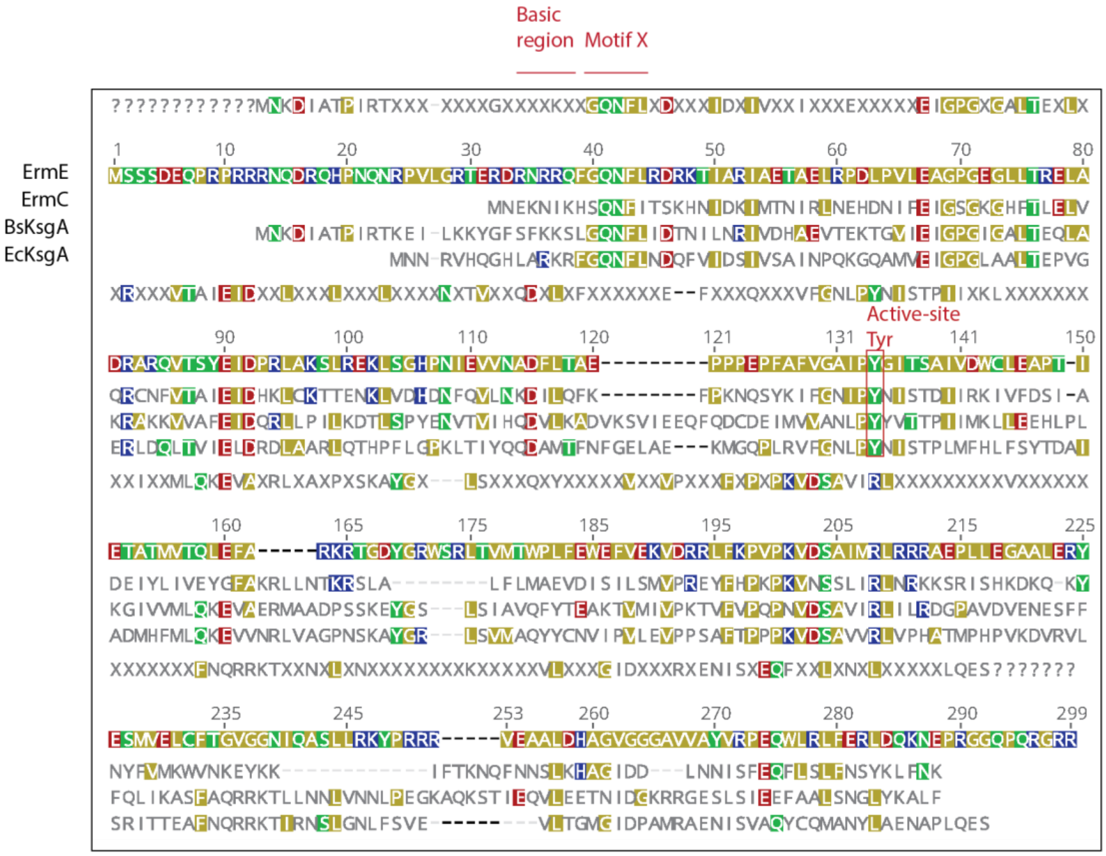


**Figure S3. Multiple sequence alignment.** A Clustal Omega alignment of *S. erythraea* ErmE, *B. subtilis* ErmC, *B. subtilis* KsgA and *E. coli* KsgA. The ErmE construct has a low sequence-complexity C-terminal region removed. Numbering is according to the ErmE sequence.

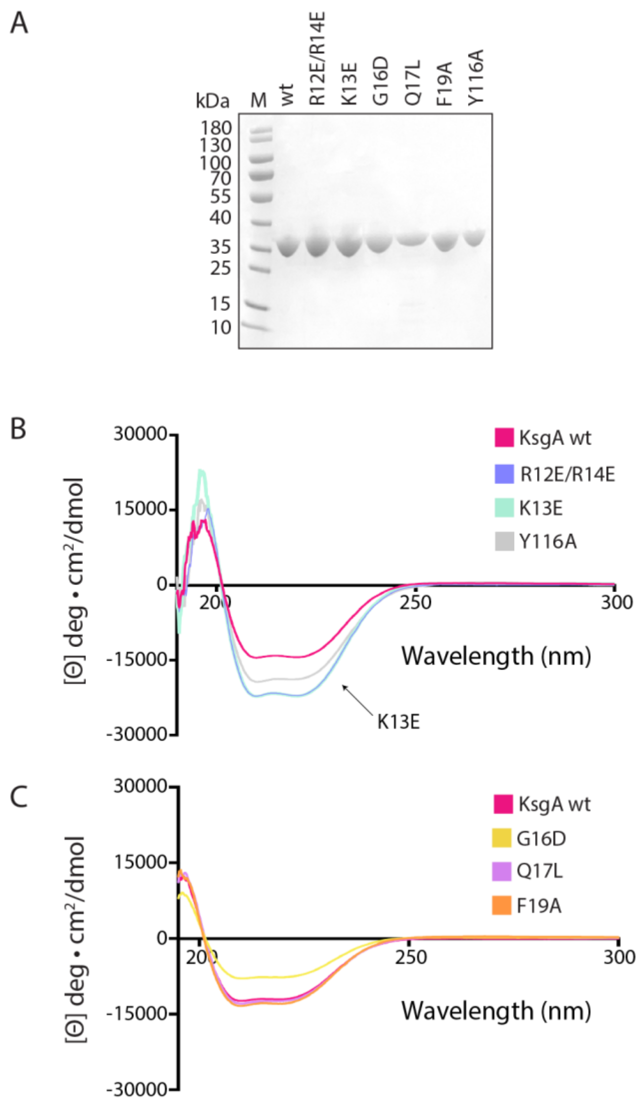


**Figure S4. Selected site-directed mutants of KsgA maintain native structure.** (A) SDS-PAGE analysis of wt KsgA and site-directed mutants is shown. (B) Circular dichroism spectroscopy of site-directed mutants within the N-terminal basic patch and motif X is shown.

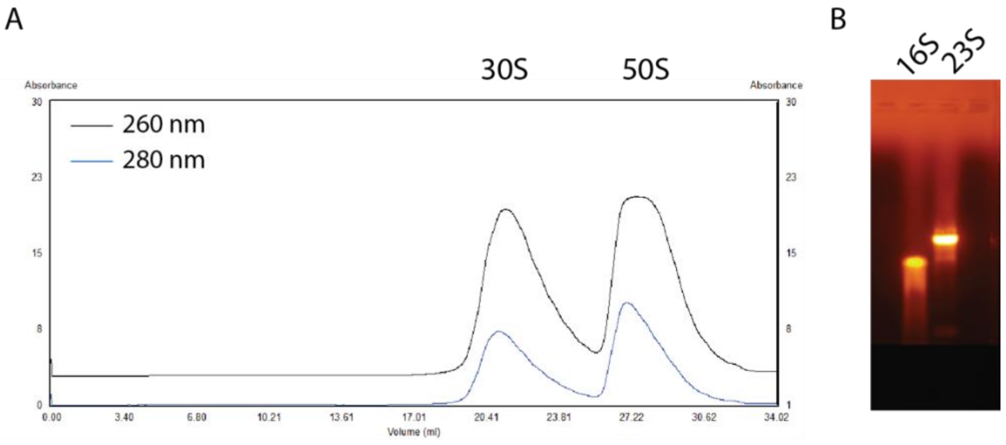


**Figure S5. Purification of 30S ribosomal subunits.** (A) Elution profile from ultracentrifugation of 10%-40% w/v sucrose gradient loaded with pelleted ribosomal subunits. (B) A 0.8% w/v agarose gel of the 30S fraction and 50S fraction from the sucrose gradient displays bands consistent with the 16S rRNA and 23S rRNA. The fractions containing 30S subunits were used as substrates for in vitro methylation by KsgA.

**Table S1. Oligos used in this study.**

| Name | Use | F/R | Sequence |
|---|---|---|---|
| prDAM001 | Genemorph pBAD *ermE* | F | GTTTTTTGGGCTAACAGGAGGAATTAAC |
| prDAM002 | Genemorph pBAD *ermE* | R | GCCCCTTCCAGCAGTGG |
| prDAM041 | *ermC* K4E Q5 | F | CATGAACGAGGAGAATATAAAACACAG |
| prDAM042 | *ermC* K4E Q5 | R | GTTAATTCCTCCTGTTAGC |
| prDAM025 | *ermC* K4E/K7E Q5 | F | ATAGAGCACAGTCAAAACTTTATTACTTC |
| prDAM026 | *ermC* K4E/K7E Q5 | R | ATTCTCCTCGTTCATGGTTAATTCC |
| prDAM015 | *ermC* S9D Q5 | F | TATAAAACACGATCAAAACTTTATTACTTCAAAACATAATATAG |
| prDAM016 | *ermC* S9D Q5 | R | TTTTTCTCGTTCATGGTTAATTC |
| prDAM017 | *ermC* Q10L Q5 | F | AAAACACAGTCTAAACTTTATTACTTC |
| prDAM018 | *ermC* Q10L Q5 | R | ATATTTTTCTCGTTCATGGTTAATTC |
| prDAM019 | *ermC* N11A Q5 | F | ACACAGTCAAGCCTTTATTACTTCAAAACATAATATAG |
| prDAM020 | *ermC* N11A Q5 | R | TTTATATTTTTCTCGTTCATGG |
| prDAM021 | *ermC* F12A Q5 | F | CAGTCAAAACGCTATTACTTCAAAACATAATATAG |
| prDAM022 | *ermC* F12A Q5 | R | TGTTTTATATTTTTCTCGTTCATG |
| prDAM023 | *ermC* I13A Q5 | F | TCAAAACTTTGCTACTTCAAAACATAATATAGATAAAATAATG |
| prDAM024 | *ermC* I13A Q5 | R | CTGTGTTTTATATTTTTCTCGTTC |
| prLOA001 | *ermC* S9D/Q10L Q5 | F | TATAAAACACGATCTAAACTTTATTACTTCAAAACATAATATAG |
| prLOA002 | *ermC* S9D/Q10L Q5 | R | TTTTTCTCGTTCATGGTTAATTC |
| prLOA003 | *ermC* S9D/N11A Q5 | F | AGCCTTTATTACTTCAAAACATAATATAGATAAAATAATG |
| prLOA004 | *ermC* S9D/N11A Q5 | R | TGATCGTGTTTTATATTTTTCTCGTTC |
| prLOA005 | *ermC* Q10L/N11A Q5 | F | AAAACACAGTCTTGCCTTTATTACTTCAAAACATAATATAG |
| prLOA006 | *ermC* Q10L/N11A Q5 | R | ATATTTTTCTCGTTCATGGTTAATTC |
| prASC011 | *ermC* Q10L/F12A Q5 | F | CGCTATTACTTCAAAACATAATATAGATAAAATAATG |
| prASC012 | *ermC* Q10L/F12A Q5 | R | TTTAGACTGTGTTTTATATTTTTCTCG |
| prLOA007 | *ermC* N11A/F12A Q5 | F | ACACAGTCAAGCCGCTATTACTTCAAAACATAATATAGATAAAATAATG |
| prLOA008 | *ermC* N11A/F12A Q5 | R | TTTATATTTTTCTCGTTCATGG |
| prDAM035 | *ermE* R34E QC | F | AGAACGTGATGAGAATCGCCGCCAATTTG |
| prDAM036 | *ermE* R34E QC | R | GTACGGCCCAACACAGGG |
| prDAM037 | *ermE* R36E QC | F | TGATCGTAATGAGCGCCAATTTGGCCAG |
| prDAM038 | *ermE* R36E QC | R | CGTTCTGTACGGCCCAAC |
| prRMP115 | *ermE* R36P QC | F | CAGAACGTGATCGTAATCCCCGCCAATTTGGCC |
| prRMP114 | *ermE* R36P QC | R | GGCCAAATTGGCGGGGATTACGATCACGTTCTG |
| prDAM039 | *ermE* R37E QC | F | TCGTAATCGCGAGCAATTTGGCCAGAACTTTCTG |
| prDAM040 | *ermE* R37E QC | R | TCACGTTCTGTACGGCCC |
| prDAM027 | *ermE* R34E/R37E Q5 | F | CGCGAGCAATTTGGCCAGAACTTTCTG |
| prDAM028 | *ermE* R34E/R37E Q5 | R | ATTCTCATCACGTTCTGTACGGCC |
| prRMP121 | *ermE* R47S QC | F | CCAGAACTTTCTGCGGGATAGCAAAACCATTGCT |
| prRMP120 | *ermE* R47S QC | R | AGCAATGGTTTTGCTATCCCGCAGAAAGTTCTGG |
| prRMP117 | *ermE* G40D QC | F | AATCGCCGCCAATTTGACCAGAACTTTCTGCGG |
| prRMP116 | *ermE* G40D QC | R | CCGCAGAAAGTTCTGGTCAAATTGGCGGCGATT |
| prASC002 | *ermE* N42A QC | F | CCGCCAATTTGGCCAGGCCTTTCTGCGGGATCGC |
| prASC001 | *ermE* N42A QC | R | GCGATCCCGCAGAAAGGCCTGGCCAAATTGGCGG |
| prACS004 | *ermE* F43A QC | F | CGCCAATTTGGCCAGAACGCTCTGCGGGATCGCAAAAC |
| prACS003 | *ermE* F43A QC | R | GTTTTGCGATCCCGCAGAGCGTTCTGGCCAAATTGGCG |
| prASC006 | *ermE* L44A QC | F | CGCCAATTTGGCCAGAACTTTGCGCGGGATCGCA |
| prASC005 | *ermE* L44A QC | R | TGCGATCCCGCGCAAAGTTCTGGCCAAATTGGCG |

| | | | |
|---|---|---|---|
| prDAM067 | *ksgA* R12E/R14E Q5 | F | AGAGTTCGGGCAAAACTTTCTCAACG |
| prDAM068 | *ksgA* R12E/R14E Q5 | R | TTCTCGGCTAAGTGGCCCTGGTG |
| prDAM079 | *ksgA* K13E QC | F | GCCCGAAGCGCTCACGGGCTAAGTGGCCCTG |
| prDAM080 | *ksgA* K13E QC | R | CAGGGCCACTTAGCCCGTGAGCGCTTCGGGC |
| prDAM081 | *ksgA* G16D QC | F | CTGATCGTTGAGAAAGTTTTGATCGAAGCGTTTACGGGCTAAG |
| prDAM082 | *ksgA* G16D QC | R | CTTAGCCCGTAAACGCTTCGATCAAAACTTTCTCAACGATCAG |
| prDAM083 | *ksgA* Q17L Q5 | F | ACGCTTCGGGCTGAACTTTCTCAAC |
| prDAM084 | *ksgA* Q17L Q5 | R | TTACGGGCTAAGTGGCCC |
| prDAM083 | *ksgA* N18A QC | F | GAACTGATCGTTGAGAAAGGCTTGCCCGAAGCGTTTACGG |
| prDAM084 | *ksgA* N18A QC | R | CCGTAAACGCTTCGGGCAAGCCTTTCTCAACGATCAGTTC |
| prDAM077 | *ksgA* F19A Q5 | F | CGGGCAAAACGCGCTCAACGATCAG |
| prDAM078 | *ksgA* F19A Q5 | R | AAGCGTTTACGGGCTAAG |
| prDAM085 | *ksgA* Y116A Q5 | F | CAACCTGCCTGCTAACATCTCCACGCCG |
| prDAM086 | *ksgA* Y116A Q5 | R | GCCGCTGCGTGTTTTCGG |
| prDAM061 | Excise *ksgA* from Top10 E. coli | F | ATGAATAATCGAGTCCACCAGGGCC |
| prDAM062 | Excise *ksgA* from Top10 E. coli | R | ACTCTCCTGCAAAGGCGCGTTCT |
| prDAM063 | Linearize pBAD IP | F | GGGCCCGAACAAAAACTC |
| prDAM064 | Linearize pBAD IP | R | GGTTAATTCCTCCTGTTAGC |
| prDAM065 | *ksgA* Homology Ends to pBAD | F | GCTAACAGGAGGAATTAACCATGAATAATCGAGTCCACCAGG |
| prDAM066 | *ksgA* Homology Ends to pBAD | R | ATGAGTTTTTGTTCGGGCCCACTCTCCTGCAAAGGCGC |

QC, QuikChange. IP, inverse PCR. F/R forward or reverse primer.

**Table S2. Gene sequences used in the study.**

| Name | Sequence |
|---|---|
| *S. erythraea ermE* (codon optimized, truncated low complexity C-terminal region) | ATGAGTAGCTCTGACGAGCAACCACGTCCACGCCGTCGCAATCAGGATC<br>GTCAACACCCGAACCAGAATCGCCCTGTGTTGGGCCGTACAGAACGTGA<br>TCGTAATCGCCGCCAATTTGGCCAGAACTTTCTGCGGGATCGCAAAACCA<br>TTGCTCGGATTGCAGAAACCGCGGAATTACGCCCGGATTTACCGGTACTG<br>GAAGCCGGACCAGGTGAAGGCCTGCTGACTCGCGAACTCGCTGATCGT<br>GCGCGTCAGGTTACGAGCTACGAGATCGATCCTCGTTTAGCCAAATCCTT<br>GCGCGAGAAACTGTCAGGCCATCCGAACATCGAGGTGGTGAATGCCGAT<br>TTCCTGACTGCCGAACCGCCGCCTGAACCGTTCGCATTCGTGGGTGCGAT<br>TCCCTACGGGATTACCAGCGCGATTGTGGACTGGTGTTTGGAGGCGCCT<br>ACCATCGAAACCGCTACCATGGTGACGCAGCTGGAGTTTGCTCGCAAAC<br>GCACGGGTGACTATGGTCGGTGGAGTCGCCTTACGGTCATGACCTGGCC<br>GCTTTTCGAATGGGAGTTCGTGGAGAAAGTGGATCGGCGCCTCTTTAAA<br>CCCGTCCCGAAAGTCGATTCGGCCATTATGCGCCTGCGTCGCCGCGCAGA<br>ACCACTGCTGGAAGGGGCAGCGCTGGAACGTTACGAATCGATGGTAGA<br>ACTGTGCTTTACAGGCGTTGGCGGCAACATCCAGGCGTCCTTACTCCGCA<br>AGTATCCCCGTCGCCGTGTTGAAGCGGCACTGGACCATGCGGGTGTTGG<br>CGGAGGGGCCGTAGTCGCCTATGTTCGCCCGGAACAGTGGCTTCGCCTG<br>TTTGAACGCCTGGACCAGAAGAACGAACCGCGTGGCGGTCAACCGCAA<br>CGTGGTCGTCGCCTCGAGCTTGGGCCC**GAACAAAAACTCATCTCAGAA**<br>**GAGGATCTGAATAGCGCCGTCGACCATCATCATCATCATCAT**TGA |
| *B. subtilis ermC'* | ATGAACGAGAAAAATATAAAACACAGTCAAAACTTTATTACTTCAAAACAT<br>AATATAGATAAAATAATGACAAATATAAGATTAAATGAACATGATAATATCT<br>TTGAAATCGGCTCAGGAAAAGGGCATTTTACCCTTGAATTAGTACAGAG<br>GTGTAATTTCGTAACTGCCATTGAAATAGACCATAAATTATGCAAAACTAC<br>AGAAAATAAACTTGTTGATCACGATAATTTCCAAGTTTTAAACAAGGATAT<br>ATTGCAGTTTAAATTTCCTAAAAACCAATCCTATAAAATATTTGGTAATATA<br>CCTTATAACATAAGTACGGATATAATACGCAAAATTGTTTTTGATAGTATAG<br>CTGATGAGATTTATTTAATCGTGGAATACGGGTTTGCTAAAAGATTATTAA<br>ATACAAAACGCTCATTGGCATTATTTTTAATGGCAGAAGTTGATATTTCTAT<br>ATTAAGTATGGTTCCAAGAGAATATTTTCATCCTAAACCTAAAGTGAATAG<br>CTCACTTATCAGATTAAATAGAAAAAAATCAAGAATATCACACAAAGATAA<br>ACAGAAGTATAATTATTTCGTTATGAAATGGGTTAACAAAGAATACAAGA<br>AAATATTTACAAAAAATCAATTTAACAATTCCTTAAAACATGCAGGAATTG<br>ACGATTTAAACAATATTAGCTTTGAACAATTCTTATCTCTTTTCAATAGCTAT<br>AAATTATTTAATAAGCTTGGGCCC**GAACAAAAACTCATCTCAGAAGAGG**<br>**ATCTGAATAGCGCCGTCGACCATCATCATCATCATCAT**TGA |
| *E. coli ksgA* | ACGATTTATCAGCAGGATGCGATGACCTTTAACTTTGGTGAACTGGCCGA<br>GAAAATGGGTCAGCCGCTGCGTGTTTTCGGCAACCTGCCTTATAACATCT<br>CCACGCCGTTGATGTTCCATCTGTTTAGCTATACTGATGCCATTGCCGACAT<br>GCACTTTATGTTGCAAAAAGAGGTGGTGAATCGTCTGGTTGCAGGACCG<br>AACAGCAAAGCGTATGGTCGATTAAGCGTCATGGCGCAATACTATTGCAA<br>TGTGATCCCGGTACTGGAAGTACCGCCGTCAGCCTTTACACCACCACCCA<br>AAGTGGATTCCGCCGTCGTGCGCCTGGTTCCTCATGCAACGATGCCTCAC<br>CCGGTTAAAGATGTTCGTGTGTTGAGCCGCATCACCACCGAAGCCTTTAA |

| | |
|---|---|
| | CCAGCGTCGTAAAACCATTCGTAACAGCCTCGGCAACCTGTTTAGCGTCG<br>AGGTGTTAACGGGAATGGGGATCGACCCGGCGATGCGAGCGGAAAATA<br>TCTCTGTCGCGCAATATTGCCAGATGGCGAACTATCTGGCGGAGAACGCG<br>CCTTTGCAGGAGAGTGGGCCC**GAACAAAAACTCATCTCAGAAGAGGAT**<br>**CTGAATAGCGCCGTCGACCATCATCATCATCATCAT**TGA |

MYC epitope and hexahistidine tag are shown in bold.
